# Benchmarking tissue- and cell type-of-origin deconvolution in cell-free transcriptomics

**DOI:** 10.64898/2026.03.05.709833

**Authors:** Alexis Ioannou, Elias T Friman, Carsten O Daub, Wendy A Bickmore, Simon C Biddie

## Abstract

Plasma cell-free RNA (cfRNA) reflects tissue- and cell-type-specific activity across pathological states and is a promising biomarker for organ injury and disease. Computational deconvolution methods are widely used to infer organ and cell-type contributions to cfRNA profiles. However, most were originally developed for single-tissue bulk transcriptomes and their performance in body-wide cfRNA settings, where any tissue or cell type can contribute, remains poorly characterised. Here, we present a systematic benchmarking of tissue- and cell type-of-origin deconvolution for plasma cfRNA that considers both methodological and reference-related sources of variability under realistic cfRNA simulation settings. We evaluated seven commonly used deconvolution methods across distinct algorithmic classes and multi-organ reference configurations derived from bulk and single-cell atlases. We assessed performance using simulation frameworks that model multi-organ mixtures, technical noise, and transcript degradation. We further examined deconvolution methods across multiple previously published clinical cfRNA cohorts spanning diverse disease contexts. Across both tissue- and cell-type-level analyses, deconvolution performance was strongly influenced by both method choice and reference parameters. Tissue-of-origin inference was comparatively robust across simulated and clinical datasets, recovering disease-associated organ signals and concordance with biochemical markers. In contrast, cell type-of-origin inference showed greater variability and reduced consistency across analytical settings, leading to divergent interpretations in both simulations and published clinical cfRNA cohorts. Together, these findings demonstrate that methodological and reference-related variability are major sources of uncertainty in cfRNA deconvolution, with tissue-level inference being more robust than cell-type-level inference. Our benchmarking framework provides guidance for reference selection and comparative interpretation in cfRNA deconvolution.

## Introduction

Plasma cell-free RNA (cfRNA) is a dynamic analyte whose composition fluctuates across physiological conditions and disease states. It contains tissue- and cell-type-specific transcripts originating from multiple organs^1,2^. Although the biological processes governing the rate and amount of cfRNA release are not fully understood, it is thought to arise from both cell death and active secretion^3,4^. cfRNA has demonstrated utility in a range of pathologies^5,6^, including malignancies^7–9^, infectious diseases^10,11^, acute conditions^12^, neurodegenerative disorders^13^, and both acute and chronic organ injury^14,15^.

The source of cfRNA can be inferred at the tissue- or cell-type level using computational deconvolution approaches^7,16^. These methods span multiple algorithmic classes, ranging from regression–based approaches to probabilistic models, but were originally developed for single-tissue bulk transcriptomes, to deconvolve into cell-types^17^, typically relying on single-tissue reference expression profiles^18^. In multi-organ disease contexts, cfRNA deconvolution has been used to identify tissues and cell types affected by pathology^7,16^. These inferred contributions were consistent with biochemical markers of organ injury, supporting the biological relevance of cfRNA deconvolution results^14^. cfRNA deconvolution has also been used to resolve multi-organ involvement in systemic disease, such as COVID-19^12^. Despite these promising applications, previous deconvolution benchmarks performed using RNA from single tissues have demonstrated variability in performance^18–22^, highlighting the need for context-specific assessment of method–reference combinations when considering cfRNA deconvolution.

Compared with single-tissue settings, body-wide cfRNA deconvolution is more challenging. Plasma cfRNA contains a larger number of potential contributors including hematopoietic and solid tissue cells, many of which exhibit overlapping transcriptional programmes^4,16,23–25^. cfRNA profiles can also be affected by molecular stability, preanalytical variability, and heterogeneous expression patterns^26,27^. In single-tissue benchmarking studies, incomplete reference profiles can lead to misassignment of signals to transcriptionally similar contributors^28–30^. Many cfRNA studies have used the human cell atlas Tabula Sapiens v1 for reference, which is an incomplete body atlas as it does not include brain cell types^12,14,16,23,31^.

A previous plasma cfRNA deconvolution benchmarking study evaluated cell-type performance using pseudo-bulk mixtures^32^. However, key challenges relevant to cfRNA deconvolution remain incompletely addressed. These include the impact of reference mismatch on cell-type inference, the accuracy of tissue-level deconvolution, and the performance of methods commonly used in cfRNA studies, such as nuSVR (nu Support Vector Regression) and BayesPrism^16,33^.

In this study, we present a comprehensive benchmarking of tissue-of-origin (TOO) and cell type-of-origin (COO) deconvolution for plasma cfRNA that explicitly considers both method and reference-related sources of variability under realistic cfRNA simulation settings. We evaluated seven common deconvolution methods: CIBERSORTx^34^, MuSiC^35^, BayesPrism^33^, ReDeconv^36^, nuSVR, quadratic programming (QP), and non-negative least squares (NNLS)^16^. These were assessed across multiple reference constructions using different simulated scenarios and validated in diverse clinical cfRNA cohorts (**Fig. 1**). By jointly examining tissue- and cell-type-level inference under matched conditions, this work provides a systematic comparison of current cfRNA deconvolution approaches and delineates how reference selection and method choice shape inferred tissue and cell-type signals in plasma cfRNA.

**Figure 1:**
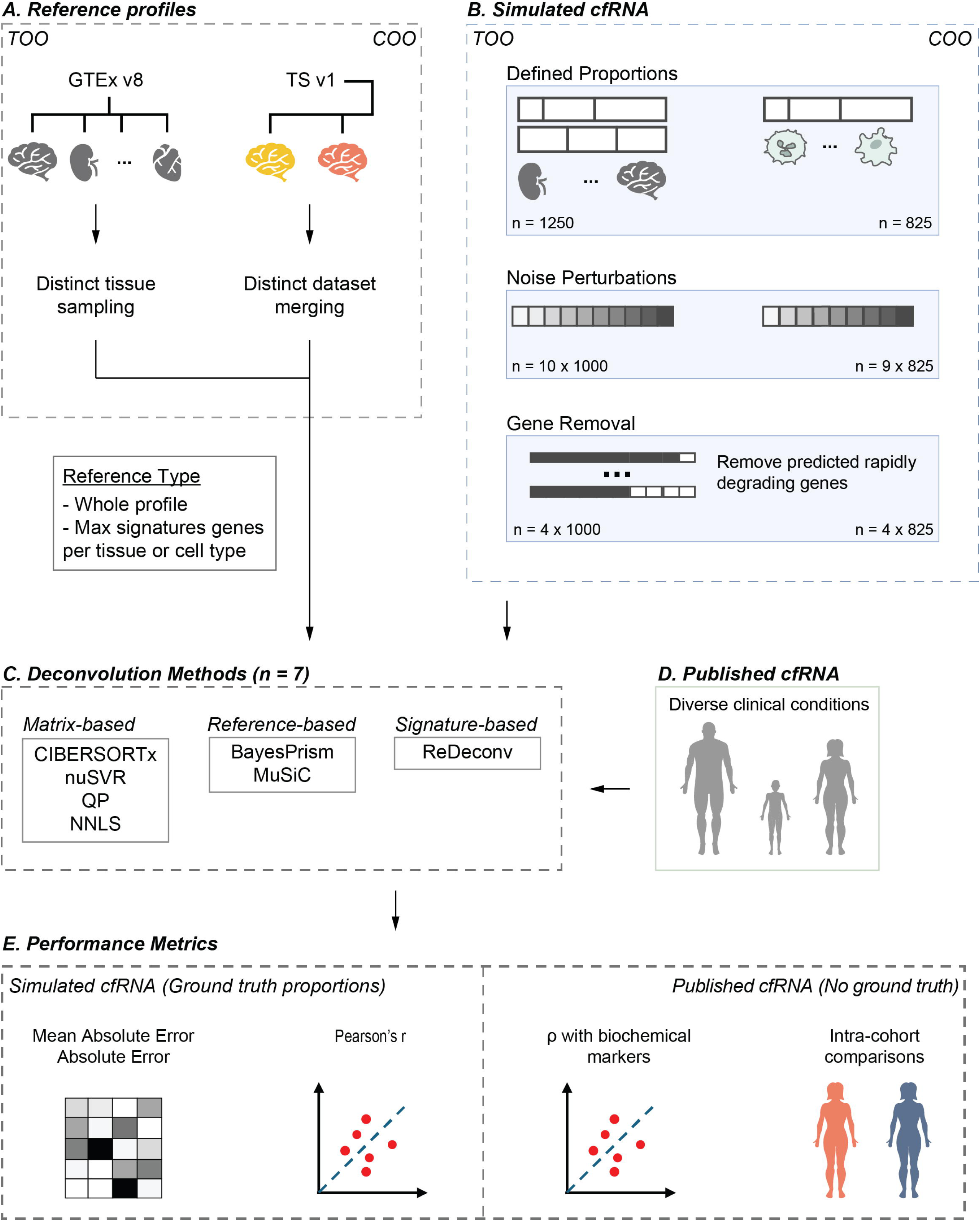
Overview of the cell-free RNA (cfRNA) deconvolution benchmarking framework for tissue-of-origin (TOO) and cell type-of-origin (COO). **(A)** Reference data were generated separately for TOO and COO deconvolution, using bulk tissue RNA-seq profiles from GTEx v8 for TOO and cell-type–resolved single-cell or single-nucleus RNA-seq datasets for COO (Tabula Sapiens v1, Darmanis brain dataset [BDa], and Human Brain Cell Atlas [HBA]). **(B)** References were assessed using simulated cfRNA generation strategies, including tissue-level mixtures (TOO) and pseudo-bulk samples derived from single-cell data (COO), with defined mixture proportions, controlled noise perturbations, and gene-removal schemes. **(C)** Seven deconvolution methods were evaluated, spanning matrix-based, signature-based, and reference-based methodological classes. **(D)** Deconvolution performance was further evaluated using published cfRNA cohorts without ground truth, across paediatric, adult, and disease contexts. **(E)** Performance was assessed using accuracy metrics for simulated data (absolute errors and correlation with ground truth) and biological validation for published datasets using biochemical markers and intra-cohort comparisons.

## Results

### Benchmarking method performance for tissue-of-origin deconvolution

To compare deconvolution methods for their ability to infer tissue-of-origin (TOO), we designed a benchmark framework based on simulated mixtures with known composition as ground truth. We evaluated seven deconvolution methods across nine reference combinations (**Fig. 1**, **Supplementary Fig. S1**). These methods were originally designed for deconvolution of bulk transcriptomes into fewer contributing cell types and represent different algorithmic frameworks (**Table 1**).

**Table 1:**
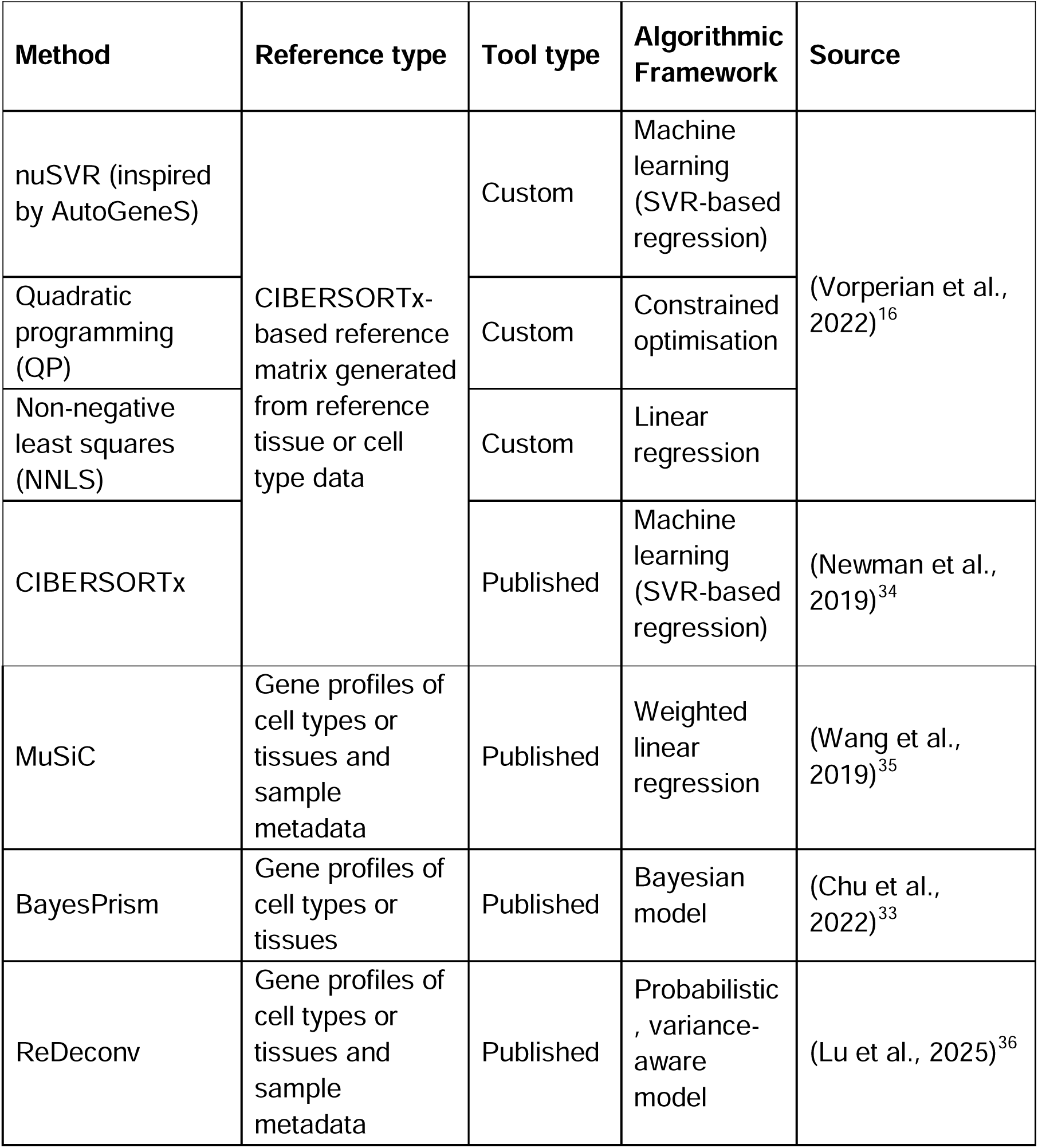
Description of the deconvolution methods assessed in this study.

Based on the required input reference, we divided methods into three categories:

1. matrix-based, which use a preconstructed matrix as their reference;
2. reference-based, which leverage full expression profiles; and
3. signature-based, which use expression profiles with a user-defined number of signatures per contributor.

Using GTEx v8 data as input reference, tissues were merged according to transcriptional similarity and shared biological function (**Fig. 2A**). Multiple reference constructions based on distinct sampling strategies (Central, Random-5, and Random-10; see Methods) were evaluated using different maximum signatures per tissue. Simulated tissue mixtures of random or uniform proportions were used, and performance was assessed using mean absolute error (MAE) and Pearson correlation between known and predicted tissue proportions.

**Figure 2:**
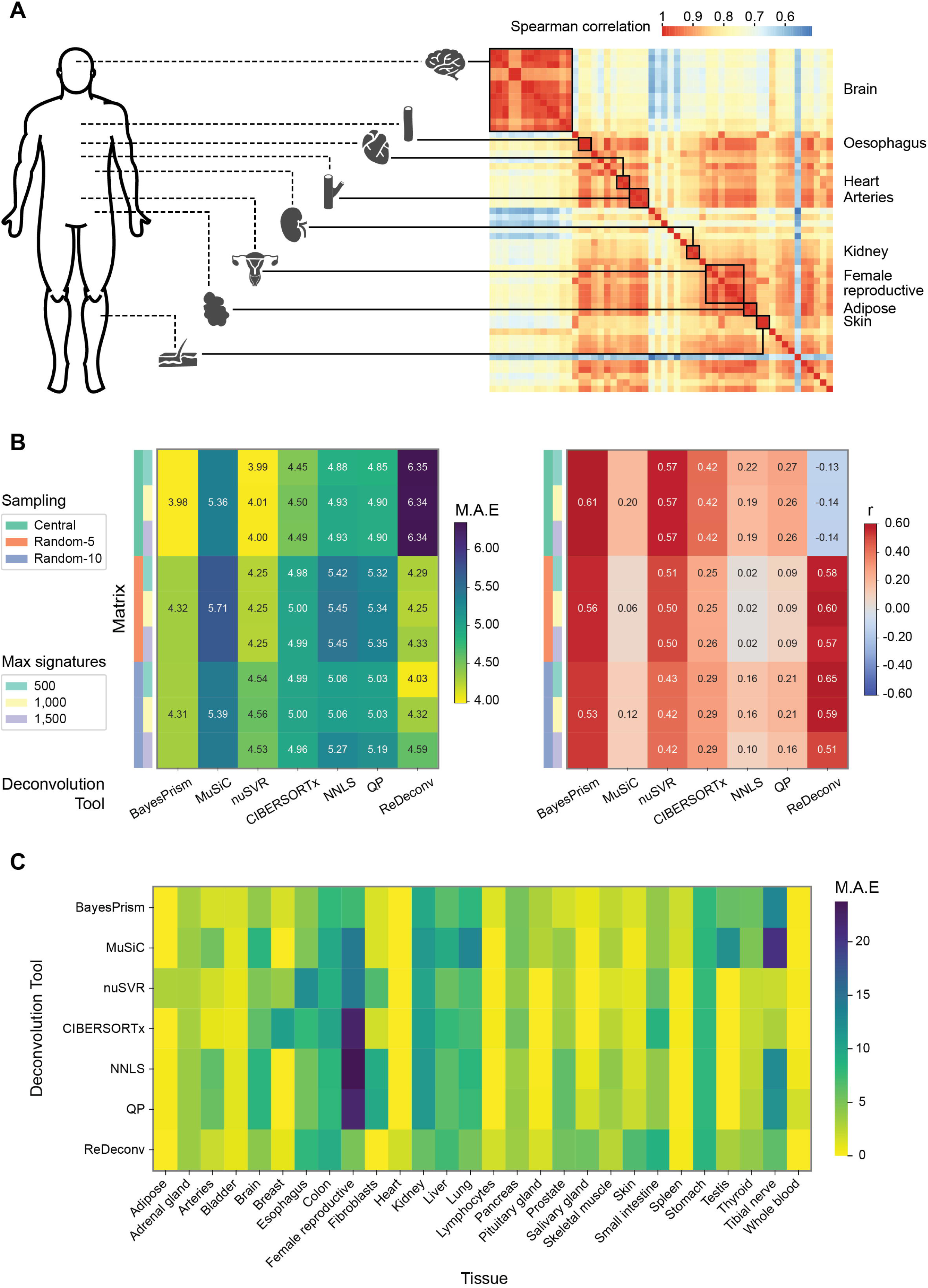
Benchmarking TOO deconvolution using 1,000 simulated cfRNA mixtures with random proportions. **(A)** Strategy for merging of GTEx v8 tissues to construct tissue reference matrices. The heatmap shows pairwise Spearman correlation (ρ) between tissues, computed using the union of the top 1,000 genes ranked by their tissue-specificity score (TSS) per tissue. Black boxes highlight tissues merged based on biological similarity and shared tissue-specific expression markers. **(B)** Performance of TOO deconvolution across seven methods and nine reference configurations, summarised as (left) heatmaps of mean absolute error (MAE, lower is better) and (right) Pearson correlation (r, higher is better) between inferred and true tissue proportions. **(C)** Heatmap showing per-tissue MAE across deconvolution methods, highlighting tissue-specific variability in deconvolution accuracy (spillover).

Across 1,000 mixtures with random tissue proportions from the same donor, deconvolution accuracy varied across tools and matrix combinations (**Fig. 2B**). BayesPrism achieved the best overall performance on the Central reference, yielding the lowest mean absolute error (MAE = 3.98) and high correlation with ground truth (r = 0.61). nuSVR achieved comparable accuracy on the same reference using a maximum of 500 signatures per tissue (MAE = 3.99; r = 0.57). ReDeconv also performed well on the Random-10 reference with 500 signatures (MAE = 4.03; r = 0.65), but consistently underestimated tissue contributions across the full range of values and yielded negative Pearson correlations under the Central sampling configuration (**Fig. 2B**; **Supplementary Fig. S2A**). MAE distributions across tools and signature set sizes are provided in **Supplementary Fig. S2B**. Analysis of 250 simulated mixtures with uniform tissue proportions yielded a similar separation between the three top-performing methods and the remaining approaches (**Supplementary Fig. S3**).

Tissue-level performance varied across methods when evaluated using their best-performing references (**Fig. 2C**). Kidney, stomach, liver, lung, the merged oesophagus group, and the merged female reproductive tissue group showed consistently higher errors across most approaches, with matrix-based methods particularly affected. In mixtures where specific tissues were absent by design (0% ground-truth proportion), non-zero contributions were frequently assigned (**Supplementary Fig. S4A**), with systematic overestimation observed for some tissues and tool-specific effects for others. Restricting MAE analysis to these absent tissues yielded a similar pattern of method performance, indicating that overestimation of absent tissues was not specific to any of the assessed methods, likely reflecting error arising from reference construction (**Supplementary Fig. S4B**).

### Robustness of tissue-of-origin deconvolution

To assess the robustness of TOO deconvolution to technical variability, we introduced increasing levels of negative-binomial noise into simulated tissue mixtures and evaluated each tool using its best-performing reference (**Fig. 3A**). BayesPrism and ReDeconv maintained similar performance across noise levels, as reflected by stable MAE values and Pearson correlations, whereas all other methods showed progressive increases in error and reduced correlation with increasing noise. Introducing noise did not substantially alter tissue-level performance patterns for most tissues (**Supplementary Fig. S5**).

**Figure 3:**
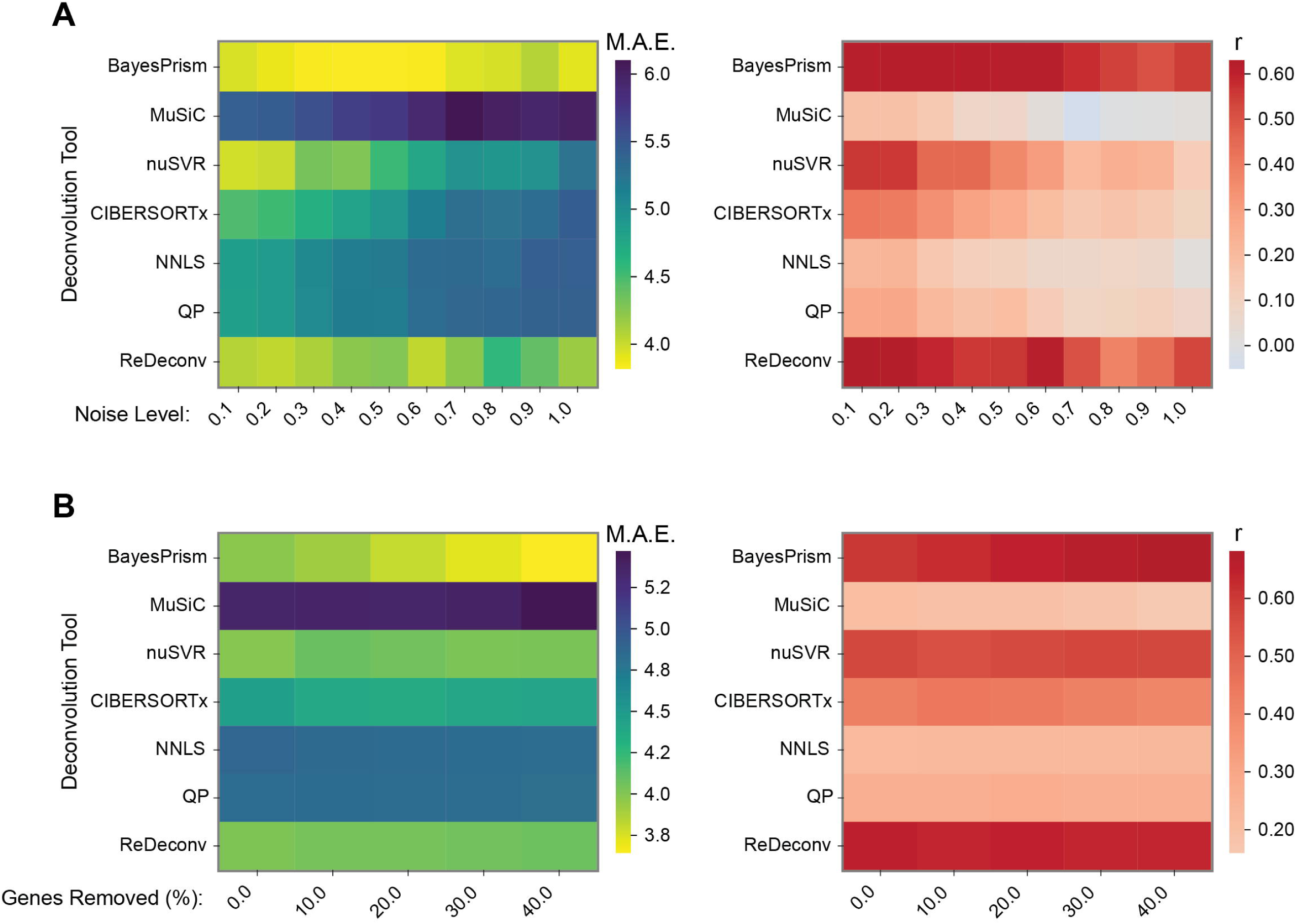
Robustness of TOO deconvolution to noise and transcript loss. (**A**) Performance of TOO deconvolution under increasing levels of negative binomial noise, evaluated using (left) mean absolute error (MAE) and (right) Pearson correlation (r). **(B)** As in **(A)** but following progressive removal of rapidly degrading transcripts in deciles. All analyses use the best-performing reference configuration for each deconvolution method.

RNA molecules have different half-lives, influenced by RNA-binding proteins and sequence features such as GC-content^37^. Stratifying genes by mRNA half-life revealed a monotonic increase in expression with transcripts stability across both GTEx tissues and cfRNA datasets (**Supplementary Fig. S6**). Given this relationship, transcript half-life may affect cfRNA deconvolution results. To examine deconvolution sensitivity to transcript instability, we sequentially removed 10 to 40% of the most rapidly degrading genes from the simulated mixtures and repeated deconvolution analyses. BayesPrism showed stable or slightly improved performance as gene removal increased, while most other methods exhibited largely stable behaviour, with only modest declines observed for MuSiC and CIBERSORTx (Fig. 3B). Overall, while technical noise marginally reduced performance, the removal of short-lived RNAs did not have a large or systematic influence on TOO deconvolution results.

### Benchmarking performance of cell-type of origin deconvolution methods

To evaluate cell type-of-origin (COO) deconvolution, we assessed the seven methods (**Table 1**) in a cell-type context. Custom reference expression profiles were generated by augmenting Tabula Sapiens v1 with human brain single-cell datasets from two independent sources^23,38,39^. Pseudo-bulk mixtures with random proportions were simulated by aggregating cells from the same donors (**Fig. 1**; **Supplementary Fig. S7**).

Across 825 simulated mixtures composed of non-brain cell types, deconvolution accuracy varied across methods and reference constructions (**Fig. 4A**). BayesPrism achieved the lowest MAE and highest correlation across reference constructions (MAE = 0.85–0.87; r = 0.82–0.85), followed by ReDeconv (MAE = 1.06–1.14; r = 0.76–0.79) and CIBERSORTx (MAE = 1.08–1.19; r = 0.73–0.82). Relative method rankings were largely preserved across reference constructions and maximum signature sizes. MAE distributions across methods, reference constructions, and signature sizes are provided in **Supplementary Fig. S8A**. Predicted versus true tissue fractions for the best-performing HBA Inner configurations are shown in **Supplementary Fig. S8B**. For MuSiC, some cell types were automatically excluded due to internal filtering, resulting in fewer predicted cell types than provided in the input reference.

**Figure 4:**
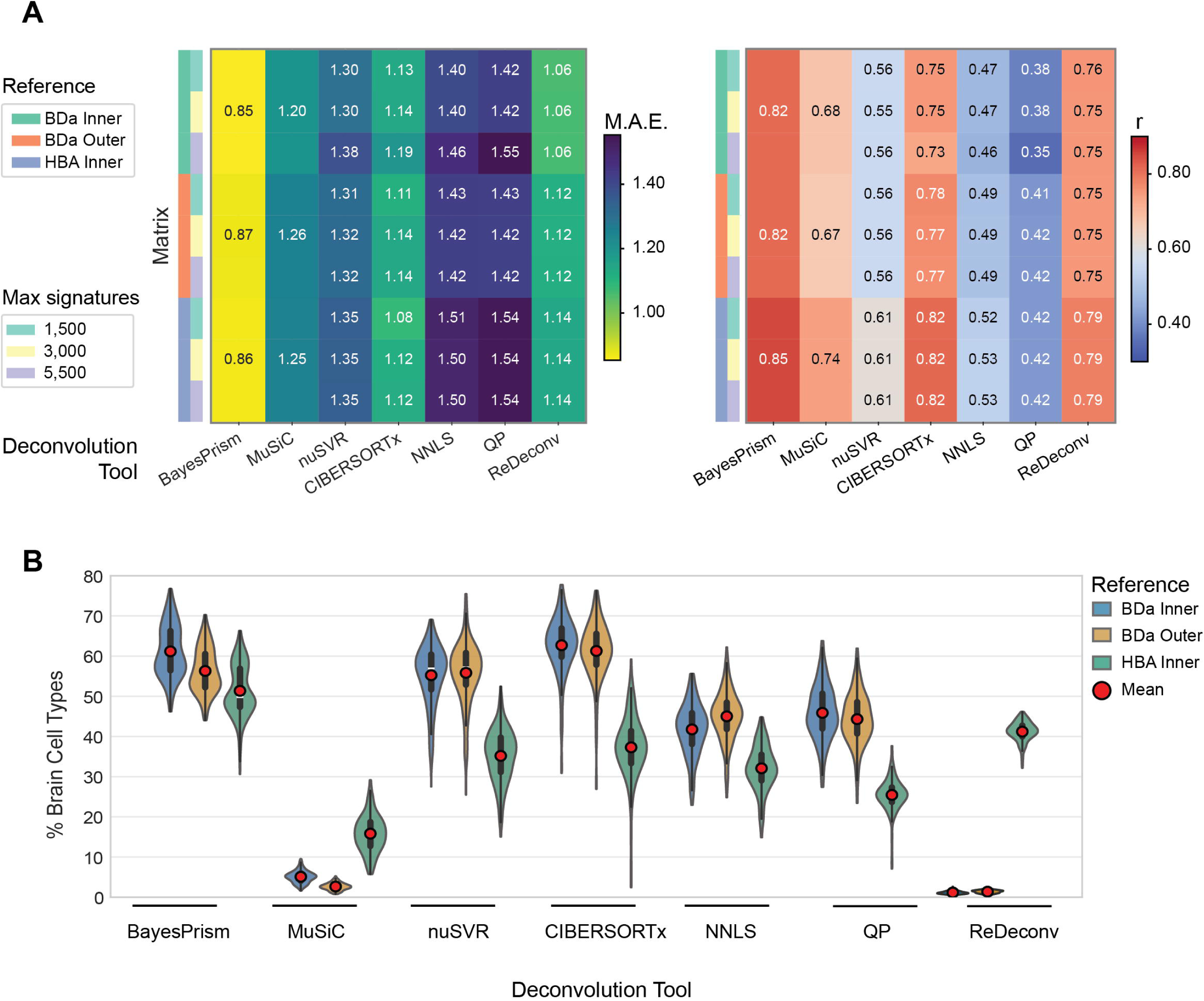
Benchmarking of COO deconvolution using 825 simulated cfRNA mixtures with random proportions. **(A)** Performance of COO deconvolution across seven methods and nine cell-type reference configurations, summarised as heatmaps of (left) mean absolute error (MAE) and (right) Pearson correlation (r) between inferred and true cell-type proportions for 825 simulated cfRNA mixtures with random proportions. Reference configurations correspond to augmentations of the Tabula Sapiens multi-organ atlas with brain single-cell data from the Darmanis brain dataset (BDa) or the Human Brain Atlas (HBA), using inner (gene intersections) or outer (gene union) gene sets, where applicable. **(B)** Distribution of inferred brain cell-type proportions obtained from deconvolution of bulk brain RNA-seq data from GTEx, using the best-performing parameter configuration per method (where applicable) across three distinct augmentations (BDa inner, BDa outer, and HBA inner).

To further characterise COO performance beyond aggregate accuracy metrics, we examined error profiles at the level of individual cell types, focusing on the 10 highest-error cell types per method. A subset of cell types consistently ranked among the highest-error contributors across reference constructions and showed elevated error across all methods (**Supplementary Fig. S9A–C**). In contrast, additional high-error cell types were specific to particular references or deconvolution methods, indicating differential sensitivity to specific cellular profiles and reference parameters rather than a single systematic artefact. Together, these results indicate that the highest COO errors are driven by a limited subset of cell types, with both shared and method-specific contributors, predominantly from immune populations. As the Tabula Sapiens v2 multi-organ reference dataset used to generate simulated COO mixtures did not include brain cell types^40^, we next evaluated deconvolution performance in a brain-only setting using 200 bulk brain samples from GTEx^41^. Each method was applied using the best-performing configuration identified in the simulation benchmark. The total proportion assigned to brain cell types varied across methods and reference augmentations (**Fig. 4B**). BayesPrism consistently assigned the highest brain cell-type proportions, while the optimal reference augmentation varied across methods, indicating sensitivity to reference composition. Notably, MuSiC and ReDeconv favoured the HBA augmentation, likely reflecting its greater donor representation. Across reference constructions, a small number of non-brain cell types were recurrently detected in more than one reference, including Schwann cells and kidney epithelial populations (**Supplementary Fig. S10A–C**).

### Robustness of deconvolution for cell type-of-origin

To assess robustness of COO deconvolution to technical noise, we introduced increasing levels of negative-binomial noise and evaluated performance using absolute error and within-method Pearson correlation (**Fig. 5A**). BayesPrism retained the lowest absolute error across all noise levels. Among the top-performing methods, ReDeconv exhibited the smallest increase in absolute error (ΔAE = 7.6) and a minimal reduction in correlation (Δr = 0.03). Methods with higher baseline error, including NNLS and QP, showed the greatest error rises with increasing noise. Taken together, our findings show that BayesPrism provides the highest absolute accuracy under noisy conditions, whereas ReDeconv exhibits the greatest stability to technical perturbations.

**Figure 5:**
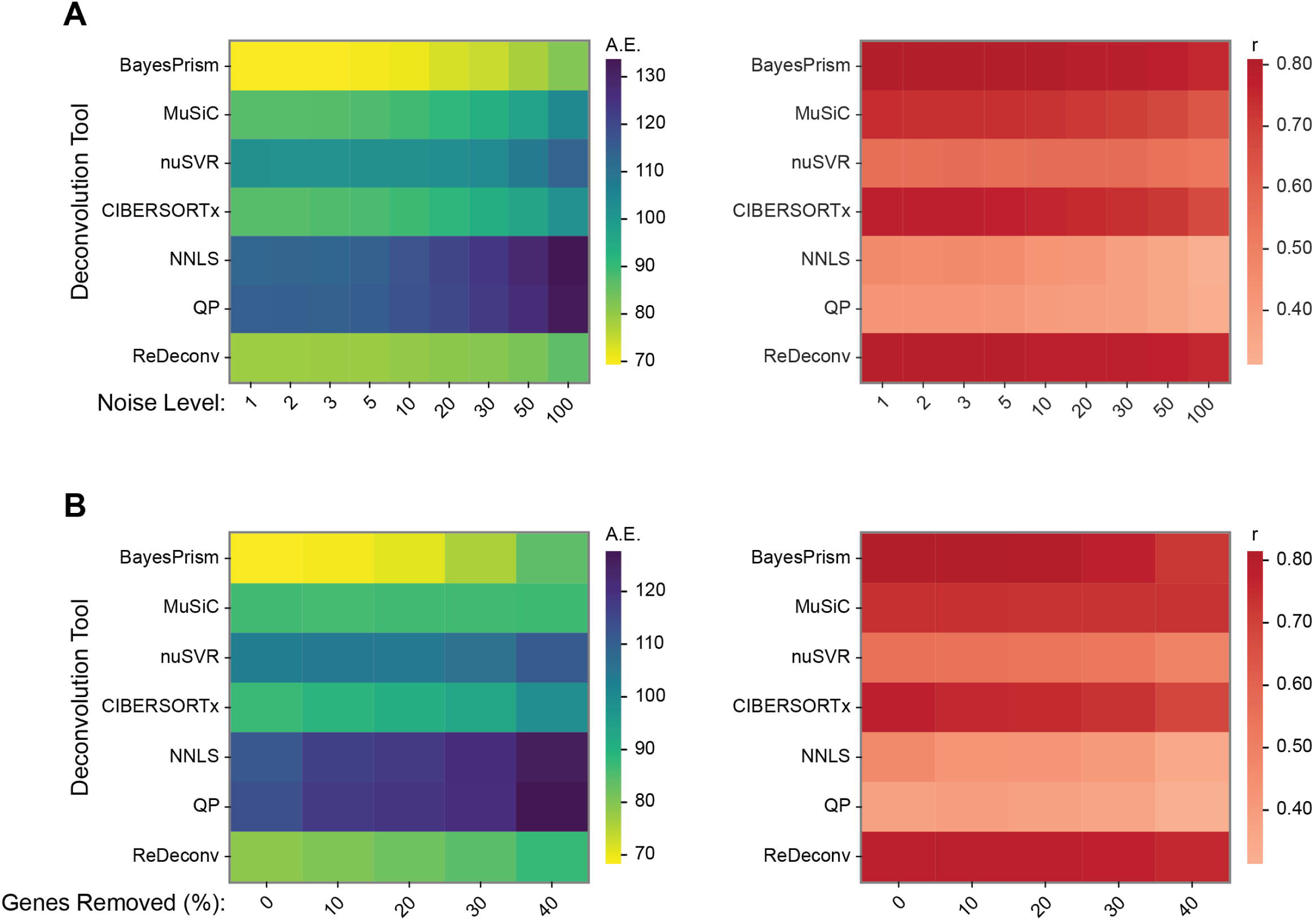
Robustness of COO deconvolution to noise and transcript loss. (**A**) Performance of COO deconvolution under increasing levels of negative binomial noise, evaluated using (left) absolute error (AE) per sample averaged across samples and (right) Pearson correlation (r). **(B)** As in (A) but following progressive removal of rapidly degrading transcripts in deciles. All analyses use the best-performing reference configuration per method.

To assess sensitivity of COO deconvolution to transcript instability, we progressively removed 10–40% of the genes with most rapidly degrading mRNAs from the simulated mixtures and repeated the deconvolution analyses (**Fig. 5B**). In contrast to the tissue-of-origin setting, degradation led to increased error across all methods. BayesPrism achieved the lowest absolute error across all levels of gene removal, followed by ReDeconv. MuSiC exhibited higher baseline error, but showed the smallest increase in absolute error (ΔAE = 1.4) and preserved correlation relative to noise-free conditions, indicating relative stability to transcript loss. Overall, technical noise and transcript loss had minimal impact on the relative ranking of COO deconvolution method, despite a reduction in accuracy.

### Tissue-of-origin deconvolution in patient-derived plasma cfRNA

To evaluate whether TOO deconvolution yields biologically meaningful signals in real cfRNA datasets and to assess the consistency of these signals across different deconvolution methods, we applied all methods to published plasma cfRNA datasets. Using the best-performing reference configuration, we analysed cohorts spanning acute injury, chronic organ disease, neurological, obstetric, and inflammatory conditions^2,12–15^. Since ground truth tissue composition is not available in these settings, we assessed performance by comparing inferred tissue contributions with biochemical markers and across disease states.

Liver injury cohorts provided a quantitative validation of TOO deconvolution. In an acute paediatric cohort with measured levels of plasma liver damage markers (alanine aminotransferase; ALT), inferred liver contribution showed a significant positive association with ALT for several methods (**Fig. 6A**; **Supplementary Fig. S11A**). BayesPrism achieved the highest Spearman correlation (ρ = 0.65), followed by CIBERSORTx and MuSiC (ρ = 0.63), and ReDeconv (ρ = 0.56). In contrast, nuSVR and QP showed weaker and non-significant associations, highlighting method-dependent variability in sensitivity to acute liver injury.

**Figure 6:**
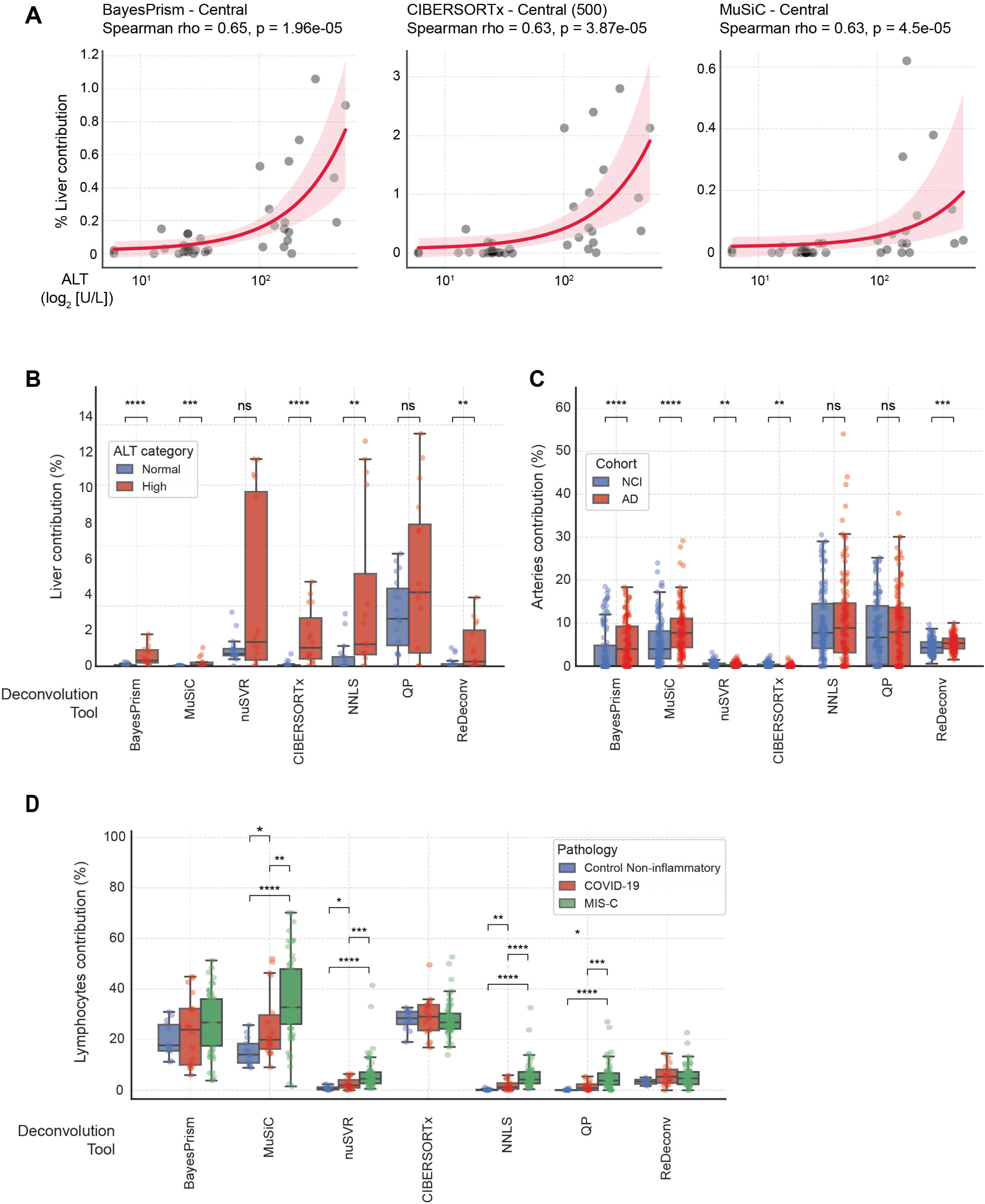
Application of TOO deconvolution in published cfRNA cohorts. **(A)** Scatterplots showing the three deconvolution methods with the highest Spearman rank correlation between inferred liver contribution and alanine aminotransferase (ALT) levels in a paediatric acute cohort; fitted trends are shown with shaded confidence intervals. The reference matrix and maximum number of signatures used for each method are indicated where relevant. **(B)** Comparison of inferred liver contribution between patients in the paediatric acute cohort stratified by ALT category (normal vs high). **(C)** Comparison of inferred arterial contributions between Alzheimer’s disease patients and non–cognitively impaired controls. **(D)** Comparison of inferred lymphocyte contribution between non-inflammatory controls, patients with COVID-19, and patients with multisystem inflammatory syndrome in children (MIS-C). Comparisons in panels **B–D** were assessed using two-sided Mann–Whitney U tests; p < 0.05 (*), p < 0.01 (**), p < 0.001 (***), and p < 0.0001 (****).

In the same cohort, stratification by previously defined ALT categories revealed significantly higher inferred liver contributions in samples with high ALT (ALT > 100 IU/L) compared with those with normal ALT (ALT < 40 IU/L) for a subset of methods (**Fig. 6B**). BayesPrism and several additional approaches detected clear separation between normal and high ALT groups, whereas other methods showed smaller or non-significant differences, with large inter-method variation in inferred liver contributions.

Similar trends were observed in a chronic liver disease cohort that contained individuals with non-alcoholic fatty liver disease (NAFLD) and non-alcoholic steatohepatitis (NASH)^15^. Inferred liver contribution correlated with both ALT and aspartate aminotransferase (AST) across methods (**Supplementary Fig. S11B**), with correlation coefficients lower than in the acute inflammatory setting. BayesPrism, CIBERSORTx, and nuSVR ranked among the top-performing methods across biomarkers and disease subsets (**Supplementary Fig. S11B**). Importantly, all methods detected significantly higher liver contributions in NAFLD/NASH than in healthy individuals (**Supplementary Fig. S12A**), despite large differences in percentage liver contribution between tools.

Beyond liver injury, TOO deconvolution revealed disease-associated tissue signals in neurological, obstetric, and inflammatory cohorts^2,12,13^, with effects that varied across methods. In Alzheimer’s disease^13^, multiple methods detected significantly increased arterial contributions relative to non-cognitively impaired controls (**Fig. 6C**). Nervous system tissues, including brain and tibial nerve, showed heterogeneous shifts between disease and control samples across methods (**Supplementary Fig. S12B-C**). In pre-eclamptic pregnancies (including severe pre-eclampsia)^2^, multiple tissues differed between normotensive and diseased groups, but both the direction and detectability of these effects varied across tools (**Supplementary Fig. S12D-F**). In inflammatory cohorts^12^, several methods inferred elevated lymphocyte contributions in COVID-19 and multisystem inflammatory syndrome in children (MIS-C; Fig. 6D). Other relevant tissues, such as tibial nerve and oesophageal mucosa, also showed increased contributions, with significance detected only by a subset of deconvolution approaches (**Supplementary Fig. S12G-H**).

Together, these results show that cfRNA deconvolution identifies biologically plausible disease-associated signals of tissue damage, but with substantial inter-method differences in inferred effect size and sensitivity.

### Application of cell type-of-origin cfRNA deconvolution to patient cohorts

Although TOO analysis captures organ-level contributions, it does not resolve the specific cellular sources underlying these cfRNA signals. To assess whether COO deconvolution yields biologically meaningful cell-type-level signals and the extent to which inference output depends on the method, we applied the COO deconvolution using the best-performing configurations to the same published cfRNA datasets as for the TOO analysis.

In the acute paediatric cohort^14^, inferred hepatocyte contributions showed a positive association with ALT for several methods (**Fig. 7A**; **Supplementary Fig. S13A**). BayesPrism achieved the strongest correlation with ALT (ρ = 0.61), followed by ReDeconv (ρ = 0.36). In contrast, MuSiC, CIBERSORTx, nuSVR, QP, and NNLS showed weaker or negative associations that did not reach statistical significance (ρ ≤ 0.31), highlighting greater divergence between methods at the cell-type level than in tissue-level analyses. Consistently, stratification by clinically defined ALT categories revealed significant hepatocyte enrichment with BayesPrism, whereas other methods showed weaker and non-significant effects (**Fig. 7B**; **Supplementary Fig. S14A**).

**Figure 7:**
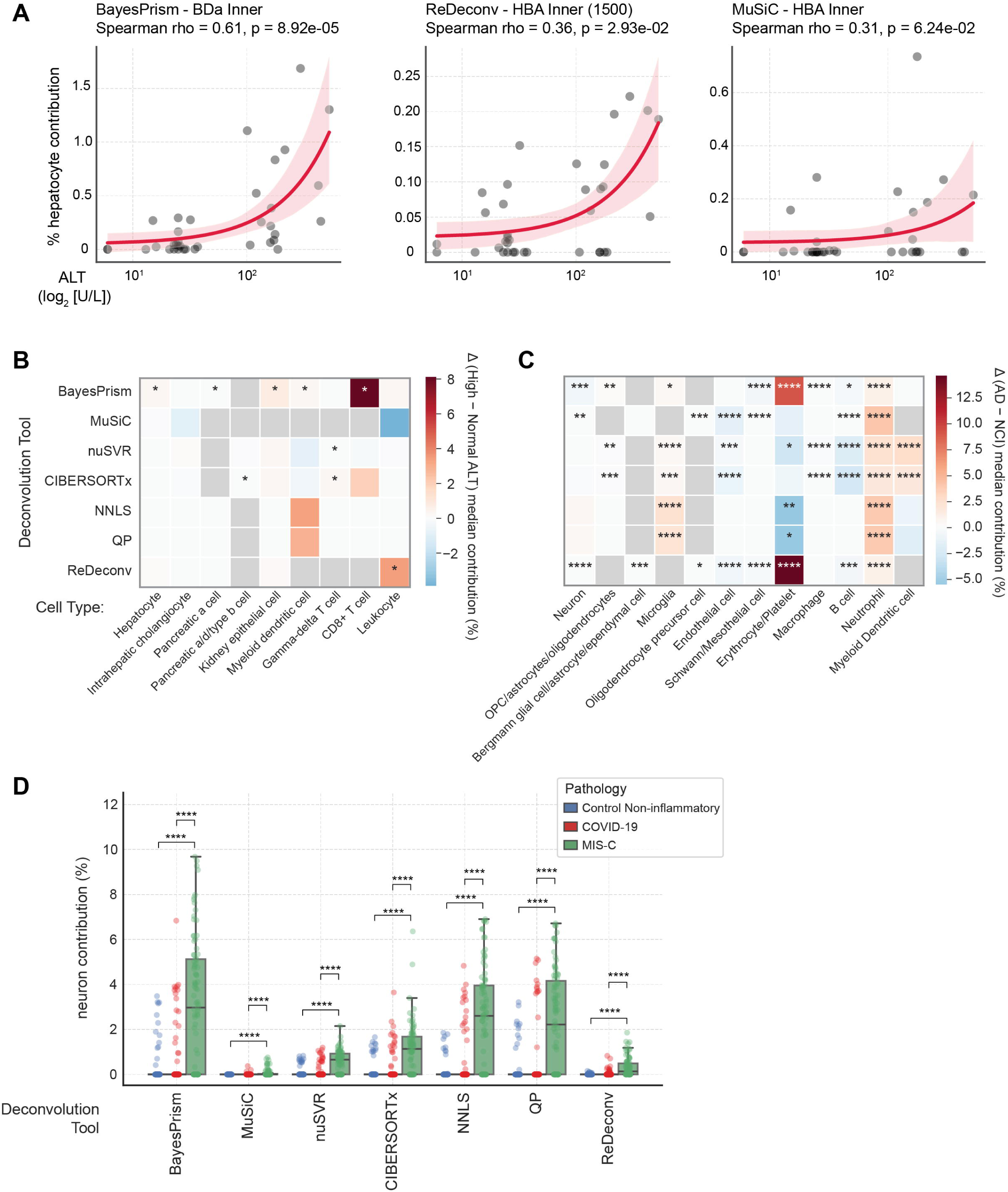
Application of COO deconvolution in published cfRNA cohorts. **(A)** Scatterplots showing the three deconvolution methods with the highest Spearman rank correlation between inferred hepatocyte contribution and alanine aminotransferase (ALT) levels in a paediatric acute cohort; fitted trends are shown with shaded confidence intervals. Where applicable, the reference matrix and maximum number of signatures used for each method are indicated. **(B)** Difference in median cell-type contributions between patients with high and normal ALT levels in a paediatric acute cohort, shown as the median contribution in the high-ALT group minus the median contribution in the normal-ALT group for each deconvolution method and cell type. **(C)** Difference in median cell-type contributions between Alzheimer’s disease patients and non–cognitively impaired controls, shown as the median contribution in Alzheimer’s disease patients minus that in non–cognitively impaired controls for each deconvolution method and cell type. **(D)** Comparison of inferred neuron contribution between non-inflammatory controls, patients with COVID-19, and patients with multisystem inflammatory syndrome in children (MIS-C). In panels B and C, grey boxes indicate cell-type categories absent from the reference matrix used by a given deconvolution method. Comparisons in panels **B–D** were assessed using two-sided Mann–Whitney U tests with Benjamini–Hochberg false discovery rate correction applied within each deconvolution method; p < 0.05 (*), p < 0.01 (**), p < 0.001 (***), and p < 0.0001 (****).

In the chronic liver disease cohort^15^, hepatocyte contributions were associated with ALT and AST across disease subsets (**Supplementary Fig. S13B**). BayesPrism and ReDeconv showed the strongest correlations across NAFLD and NASH, while MuSiC, nuSVR, and CIBERSORTx exhibited moderate, cohort-dependent performance. In contrast to the acute cohorts, multiple methods detected significantly higher hepatocyte contributions in diseased compared with healthy individuals; however, the size of these differences and the accompanying shifts in additional cell types differed across tools (**Supplementary Fig. S15A**, **S14B**).

Beyond liver diseases, COO deconvolution showed method-dependent variability across multiple diseased cohorts compared to healthy controls, with differences in both effect size and statistical support across tools. In Alzheimer’s disease^13^, some methods detected significant changes in multiple brain-associated and blood cell types (**Fig. 7C**; **Supplementary Fig. S14C-S14D**). Similarly, several cell types differed between normotensive and pre-eclamptic pregnancies (including severe cases)^2^, but both the identity of significant cell types and the direction of change varied widely across methods, with some tools detecting broad shifts and others showing only sparse or non-significant effects (**Supplementary Fig. S15B, S14E-S14F**). In inflammatory cohorts comparing COVID-19, MIS-C (multisystem inflammatory syndrome in children), and non-inflammatory controls^12^, all methods inferred elevated neuronal and immune contributions, but with variable magnitudes and statistical support (**Fig. 7D**; **Supplementary Fig. S14G-S14H**).

Collectively, these results indicate that COO deconvolution can recover plausible disease-associated cell-type signals across diverse clinical contexts, but with limited concordance across methods in both the identity and magnitude of inferred changes.

## Discussion

Plasma cell-free RNA (cfRNA) provides a body-wide readout of transcriptional activity that can be deconvolved into tissue- and cell-type-specific contributions^1,2^, making it a promising biomarker for various diseases. However, the robustness of inferred signals from computational deconvolution across analytical choices has remained poorly characterised in cfRNA settings. In this study, we performed a systematic evaluation of TOO and COO deconvolution for the cell-free transcriptome using controlled simulations and diverse clinical cfRNA datasets, providing a joint benchmark for origin inference. By benchmarking seven methods across multiple reference constructions, we show that cfRNA deconvolution accuracy is sensitive to the specific method–reference pairing, and that methods differ in robustness to noise, transcript degradation, as well as in the magnitude and statistical support of disease-associated signals. Our findings indicate that methodological variability is a major source of uncertainty in cfRNA-based inference.

At the tissue level, BayesPrism, nuSVR, and ReDeconv showed the strongest overall performance in simulations when evaluated using their best-performing reference configurations, with BayesPrism providing the most consistent balance of accuracy and stability. Similarly, BayesPrism showed the strongest concordance between inferred liver contributions and biochemical markers in liver injury cohorts, but several other methods, including CIBERSORTx, MuSiC, and ReDeconv, also exhibited comparable associations. Across cohorts, the absolute magnitude of inferred tissue fractions differed widely across methods, even when the direction and significance were consistent. This indicates that while cfRNA reliably identifies which organs contribute RNA in disease, quantitative tissue proportions remain highly method dependent and should be interpreted comparatively rather than as precise measurements.

Cell type-of-origin deconvolution exhibited greater methodological dependence than tissue-of-origin. In simulations, BayesPrism, CIBERSORTx, and ReDeconv showed the strongest overall performance, with BayesPrism and ReDeconv also demonstrating the greatest stability to perturbations. Consistent with their reliance on fine-grained transcript signatures^26^, COO methods were more sensitive to simulated transcript degradation and showed increased spillover to closely related cell types when deconvolving bulk brain data. In acute liver injury, only BayesPrism robustly detected hepatocyte contributions across both correlation with ALT and disease stratification. In other cohorts, different methods implicated different cell types and produced divergent disease-associated cellular signatures. This likely reflects the higher transcriptional similarity and larger number of potential categories at the cell-type level, which increase collinearity and reduce identifiability.

Reference data quality and completeness are well recognised challenges in deconvolution^18,26^, and their impact has been studied in immune and stromal contexts^28^. For cfRNA, we found that how tissues or cell types were sampled, merged, or augmented (such as inner vs outer merges or the use of Darmanis vs HBA brain references) in some cases altered proportions to a degree comparable to switching between deconvolution methods. Beyond these construction effects, reference incompleteness introduces a further source of bias. Most cfRNA deconvolution studies rely on Tabula Sapiens v1 as a multi-organ reference despite its lack of brain cell types^5,23^. One notable example is the deconvolution of children with COVID-19 and MIS-C, where significant increases in Schwann cells were reported^12^. The same cell type was previously identified as the largest contributor when deconvolving bulk GTEx brain samples using this reference^16^. In our analyses, augmenting Tabula Sapiens with brain datasets shifted this interpretation, with neurons showing statistically significant differences across all methods while Schwann cells were detected by only a subset of tools, potentially acting as a surrogate for missing neuronal populations.

Our results also help to contextualise a recent pseudo-bulk benchmark of the circulating transcriptome. In that study, pseudo-bulk mixtures were generated and deconvolved using the same version of Tabula Sapiens^32^, which necessarily reduces the impact of reference mismatch but introduces circularity. Here, we observed substantial reference-associated variability under mismatched conditions. This suggests that matched-reference benchmarking designs may overestimate real-world performance.

Several limitations should be acknowledged. Owing to the lack of ground truth in cfRNA datasets^26^, we relied heavily on simulations to model multi-organ mixtures. These mixtures were derived from intracellular RNA and therefore may not fully capture the fragmentation, degradation, and coverage biases characteristic of cfRNA in plasma^7,27,42^. We partially addressed this limitation by validating deconvolution signals against biochemical markers of organ injury. In addition, we lacked *in vitro*–generated multi-organ cfRNA mixtures; comparable experimental mixtures have so far been developed for immunological and immuno-oncological benchmarking^20,22^. We were also unable to model the dominance of blood-derived RNA at the tissue level because our simulations were based on multi-organ per-donor atlases that did not include paired blood samples, and therefore focused on multi-organ tissue mixtures. Finally, limited availability of normal multi-organ human datasets from the same donors required us to use a single dataset per level, particularly for TOO, which we partially mitigated by introducing noise to generate technical replicates. Although our analyses focused on plasma cfRNA, the benchmarking framework can be applicable to cfRNA from other biofluids, such as urine or cerebrospinal fluid (CSF)^43–45^, where tissue- and cell type-of-origin inference is likely to be similarly constrained by reference and methods. However, given that urinary and CSF cfRNA exhibits a distinct landscape of cell-type contributions compared with plasma^46–48^, application of these approaches to other biofluids will require biofluid-specific and organ-focused reference datasets.

As more comprehensive multi-organ, large-scale single-cell atlases become available^5,49^, including disease-specific atlases, cfRNA deconvolution will become more accurate and interpretable. Our results suggest that improvements in reference completeness, particularly for neurological and stress-associated cell populations, will be as important as advances in method design. Explicit modelling of cfRNA-specific transcript degradation and fragmentation is likely to further improve robustness, especially for cell-type-level inference. Together, these principles provide a framework for more reliable application of cfRNA deconvolution.

## Materials and Methods

### Deconvolution methods

We selected seven deconvolution tools for both tissue- and cell type-of-origin analyses, including custom implementations based on published frameworks and publicly available tools. These tools span regression-based and probabilistic approaches and were chosen based on their performance in previous single-tissue benchmarks^19,21^ and their use in prior liquid biopsy studies^12,16^. An overview of the methods and their required reference types is provided in **Table 1**.

For nuSVR, QP and NNLS, no expression level filtering of mixture samples was performed prior to scaling and deconvolution. The nuSVR algorithm was executed with a maximum runtime of 120 hours per hyperparameter combination^16^. Additionally, ReDeconv was applied without its internal normalisation step due to limited availability of matched same-sample reference profiles^36^.

### Tissue-of-origin reference generation from bulk tissue expression profiles

Gene expression matrices of human tissues were obtained from GTEx v8 RNA-seq data^41^, comprising 54 distinct tissues, which were merged into 30 tissue categories for downstream analyses. Tissue merging was based on shared biological function and similarity of tissue-specific gene expression signatures, defined using the previously described tissue-specificity score^7^ (TSS; **Supplementary Table S1**; **Fig. 2A**). Briefly, for each gene j, median expression xij across samples within tissue i was computed, and TSS was defined as:

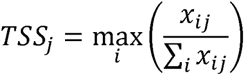

Higher TSS values indicate greater tissue specificity. For each tissue, the top 1000 genes ranked by TSS were used to assess tissue similarity and inform tissue grouping.

Three distinct reference parameters were generated for each tissue category and used as inputs for the deconvolution benchmarking analyses. The Central reference consisted of two samples per tissue whose overall gene expression profiles were closest to the group median. The Random-5 and Random-10 references were created by randomly subsampling five or ten tissue profiles, respectively, from each category (**Supplementary Fig. S1**).

For methods requiring a CIBERSORTx-based signature matrix (**Table 1**), each reference variant was processed using the command-line version of CIBERSORTx (v1.0; fractions module). Signature matrices were constructed with gene sets of 300–500, 500–1000, and 1000–1500 genes, with a q-value cutoff of 0.01. Similarly, ReDeconv was run using a maximum of 500, 1000, or 1500 signature genes. Gene identifiers were harmonised between reference profiles and mixtures prior to deconvolution. For methods requiring metadata, each tissue profile was treated as an independent specimen.

### Generation of simulated bulk samples from tissue expression profiles

RNA-seq data of bulk normal human tissue^50^ were aligned to the GRCh38 human reference genome using STAR (v2.7.8a). Non-chromosomal reads and duplicates were removed with samtools (v1.16), and gene-level expression was quantified using FeatureCounts (v2.0.1) with GENCODE v46 gene annotations. We selected donors with multiple tissue samples available, including FFPE-2, FFPE-3, FFPE-4, FFPE-5, Later-2, Later-8, Later-12, and Later-16, to enable within-donor tissue mixing. Expression matrices were normalized to counts per million (CPM) and used to generate simulated bulk samples using two approaches. In the uniform-mixing approach, 250 samples were created by averaging CPM-normalized expression values from at least three tissues per donor in equal proportions. In the random-proportion approach, 1000 samples were generated by combining four or more tissues per donor, where expression values were multiplied by randomly assigned proportions (5–85%) that summed to 100%.

### Cell type-of-origin reference generation from single-cell RNA-sequencing data

Single-cell RNA sequencing (scRNA-seq) data were obtained from the Tabula Sapiens atlas version 1 (TSP)^40^, the Human Brain Cell Atlas (HBA v1.0)^39^, and the Darmanis brain dataset (BDa)^38^. For TSP, only 10x Genomics data were retained and counts from the DecontXcounts layer were used. The HBA neuron and non-neuron datasets were merged and subsampled to 200,000 cells. For all datasets, doublets as identified by Scanpy (sc.pp.scrublet) were removed. As previously described, quality-control criteria were applied^40^, by excluding cells with fewer than 200 detected non-mitochondrial genes or fewer than 2,500 non-mitochondrial counts. TSP and BDa (excluding fetal and unpurified populations) were merged using either an inner gene set, defined as the intersection of genes shared across datasets, or an outer gene set, defined as the union of all detected genes. HBA datasets were merged using the inner gene set only (**Supplementary Fig. S7**). All preprocessing and quality-control analyses were performed using Scanpy (v1.10.4)^51^.

For each of the three augmented datasets, fine-grained cell ontology annotations were filtered and merged into broader cell-type groups using a previously defined hierarchical clustering–based approach with minor modifications (Vorperian et al., 2022). Briefly, our filtering retained a broader set of muscle cell annotations than in the original workflow and we applied a less aggressive collapsing of immune cell subtypes (**Supplementary Table S2**), allowing finer-grained immune annotations to be preserved while excluding selected broad or ambiguous immune labels. Following filtering, clustering was performed across all compartments simultaneously, and dendrograms were cut at a fixed relative height of 7.5% of the maximum dendrogram height to define merged cell-type groups.

Following cell-type merging, up to 100 cells per fine-grained cell type were randomly selected as reference for BayesPrism, MuSiC, and ReDeconv deconvolution analyses, whereas up to 25 cells per fine-grained cell type were retained for all remaining methods (**Table 1**). Sampled cells were subsequently relabelled by mapping their original fine-grained annotations to the corresponding merged cell-type groups, yielding reference matrices with columns representing merged cell types. For deconvolution methods requiring a CIBERSORTx-derived signature matrix (**Table 1**), reference matrices were generated locally using the CIBERSORTx fractions module (v1.0). Signature matrices were constructed using multiple gene set sizes (300–1500, 1000–3000, and 3000–5000 genes), with an average expression threshold of 0.25 and 10 replicates per cell type. ReDeconv analyses were performed using a maximum of 1500, 3000, or 5000 signature genes, requiring a minimum of two cells per cell type and allowing up to 15 cell types to fall below the expression threshold. For methods requiring metadata as input (**Table 1**), we used donor, tissue and sampling site information for TSP data, whereas donor and tissue information were used for the augmented brain datasets.

### Generation of simulated pseudo-bulk samples from single-cell RNA-sequencing data

To avoid circularity with reference profiles derived from earlier dataset releases, scRNA-seq data from the Tabula Sapiens v2 dataset were used to generate simulated mixtures^40^. Cells from donors TSP21, TSP25 and TSP27 profiled using the 10x Genomics platform were extracted, and donor–tissue subsets were processed individually. Cells with fewer than 100 detected genes and genes expressed in fewer than three cells were removed. Quality control followed the procedures described above. The DecontXcounts layer provided in the Tabula Sapiens v2 dataset was used as the expression matrix for all subsequent steps. To ensure compatibility with downstream deconvolution, each QC-filtered dataset per donor was restricted to cell types represented in the reference profiles.

Pseudo-bulk RNA-seq samples were generated by randomly subsampling cells from the filtered donor-level datasets. For each pseudo-bulk, between 4 and 15 eligible cell types were selected at random. The minimum number of cells required per cell type was set to 5% of the target total number of cells. Cell-type fractions were sampled from a Dirichlet distribution, and approximately 200, 400, 600, 800 or 1000 cells were drawn without replacement according to these fractions. Due to minimum per–cell-type constraints and donor-specific abundances, the final total number of cells per sample varied slightly around the target. For each pseudo-bulk, decontaminated counts were normalised per cell, summed across all selected cells, and converted to CPM to generate bulk-like expression profiles. Ground-truth cell-type compositions, defined by the exact number of contributing cells per type, were retained for benchmarking deconvolution performance.

### Assessment of evaluation performance

To evaluate the performance of TOO and COO deconvolution, absolute error (AE), mean absolute error (MAE), and the Pearson correlation coefficient (r) were used as performance metrics, calculated as follows:

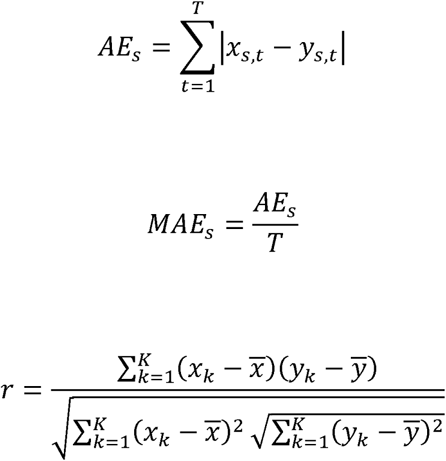

where x_s,t_ and y_s,t_ denote the estimated and ground-truth proportions, respectively, for tissue or cell-type category *t* in sample *s*, and *T* is the total number of categories. For the Pearson correlation coefficient, *k* indexes each paired predicted and ground-truth proportion across all samples and categories, with *K* denoting the total number of such pairs. Pearson correlation (r) was calculated between predicted and ground-truth proportions across all samples and categories, excluding category–sample pairs where both values were zero. MAE was computed both globally across all tissue–sample pairs and at the category level (per-category MAE, reflecting spillover between tissues or cell types). Prior to metric calculation, deconvolution outputs were harmonised to a common set of tissue/cell-type categories (including merging equivalent labels and assigning zero to unreported categories) and normalised to sum to 100% per sample. AE was used for COO comparisons involving reference sets that produced differing numbers of output categories.

### Noise robustness

Both simulated bulk (TOO) and pseudo-bulk (COO) datasets were perturbed to introduce controlled count-level noise, modelling stochastic variability in cfRNA count measurements.

For simulated bulk samples, ten linearly spaced levels of noise were added to the counts of individual tissues using a previously described negative binomial model with minor modifications^21^. Specifically, the biological coefficient of variation (BCV) scaling factor was increased from 1.8 to 3.24, yielding:

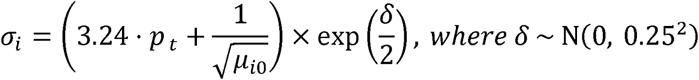

Five replicate simulations were generated per tissue per patient.

For simulated pseudo-bulk (COO) samples, noise was introduced at the single-cell level using a negative binomial (gamma–Poisson) model. Following per-cell normalization, expression values were perturbed using gene-wise overdispersion controlled by φ, applied across nine increasing noise levels (φ = 1, 2, 3, 5, 10, 20, 30, 50, 100). In both cases, the resulting datasets were combined using their corresponding mixing approach and the same tissue or cell type proportions as described above.

### Degradation analysis

Differential rates of mRNA decay are likely to contribute to variability. Therefore, human mRNA half-life data^37^ were used to assign genes into five stability categories relative to the mean (μ) and standard deviation (σ) of mRNA half-life. These were: Very Low (≤ μ - σ), Low (μ - σ to μ - 0.5σ), Medium (μ ± 0.5σ), High (μ + 0.5σ to μ + σ), Very High (≥ μ + σ). GTEx v8 read counts per tissue and counts from published cfRNA datasets^2,13–15^ were normalised to CPM. Published cfRNA sequencing data were obtained either as raw sequencing reads^13,15^ or as author-provided expression matrices^2,14^. For Moufarrej et al. dataset, these analyses used the Stanford and Validation2 cohorts using the preQC count matrices. Where raw sequencing data were available, reads were processed using the same alignment, filtering, and quantification workflow applied to bulk tissue RNA-seq data using STAR (v2.7.11b) and samtools (v1.20). Mean log_2_(CPM+1) expression was then compared across stability categories.

To assess the effect of transcript stability on deconvolution, genes were ranked by mRNA degradation rate^37^ and the top 10–40% fastest-degrading genes removed. Due to the different input gene sets, the genes removed at each percentile differed between the simulated bulk and pseudo-bulk datasets. Expression values for removed genes were set to zero and the matrices were re-normalised to CPM and subjected to deconvolution with the best-performing reference for each tool.

### Deconvolution of published cfRNA datasets

All expression matrices from published cfRNA datasets^2,12–15^ were normalised to CPM and applied to both TOO and COO frameworks, using the best-performing reference for each deconvolution tool, with gene identifiers harmonised between reference profiles and cfRNA expression matrices. For cohorts providing raw sequencing data^13,15^, the sample with the highest sequencing depth per patient was retained. For the Moufarrej cohort^2^, pre-QC count matrices were used, restricted to samples present in the corresponding post-QC metadata. For the paediatric inflammatory cohort^14^, analyses were limited to infection categories represented by more than three patients and to sepsis cases.

Deconvolution outputs were compared with biochemical blood test measurements and clinical cohort pathology data. Associations between estimated liver or hepatocyte contributions and biochemical markers of liver injury (ALT or AST) were assessed using Spearman rank correlation. Comparisons between patient groups within cohorts were performed using two-sided Mann–Whitney U tests per deconvolution tool, with patient stratification (e.g. ALT categories) defined according to the original study-specific criteria. For COO analyses, p-values were adjusted within each tool using the Benjamini–Hochberg false discovery rate correction (adjusted *P*-value < 0.05)

## Acknowledgments

We thank the members of the Bickmore and Daub Labs, Martin Taylor, and Ava Khamseh for helpful discussions, valuable input and feedback. We are also grateful to N. Chalasani and T. Maddala for sharing biochemical data for the chronic liver disease cohort. We acknowledge the Edinburgh Compute and Data Facility (ECDF) (www.ecdf.ed.ac.uk/) and the Centre for Bioinformatics and Biostatistics (CBB) at Karolinska Institutet for providing computational resources.

## Author contributions

A.I. contributed to methodology, performed data curation, formal analyses, and data visualisation, and wrote the original draft of the manuscript. W.A.B., and S.C.B. contributed to conceptualization and provided funding. E.T.F., C.O.D., W.A.B., and S.C.B. contributed project supervision. All authors contributed to the editing of the manuscript.

## Data and code availability

Data used in this study are publicly available from the cited studies unless otherwise stated. Raw sequencing reads from Suntsova et al. (PRJNA494560), Toden et al. (PRJNA574438), and Chalasani et al. (PRJNA701722) were downloaded from the Sequence Read Archive (SRA). Count matrices from Loy et al. (2024; GSE192902), Loy et al. (2023; GSE225223), Moufarrej et al. (GSE25055), and Darmanis et al. (GSE67835) were downloaded from the Gene Expression Omnibus (GEO). GTEx read counts per tissue (version 8) were downloaded from the GTEx Portal (https://www.gtexportal.org/home/datasets). Tabula Sapiens v1 and v2 were downloaded from the Tabula sapiens v2 Figshare repository (https://figshare.com/articles/dataset/Tabula_Sapiens_v2/27921984). Human Brain Cell Atlas (HBA) v1.0 datasets (neuronal and non-neuronal populations) were obtained from the Human Cell Atlas data portal (https://data.humancellatlas.org/hca-bio-networks/nervous-system/atlases/brain-v1-0).

Code used in this work is available on Github at www.github.com/Ioannou-A/cfRNA-decon.

## Supplementary Figure Legends

**Supplementary Figure S1:**
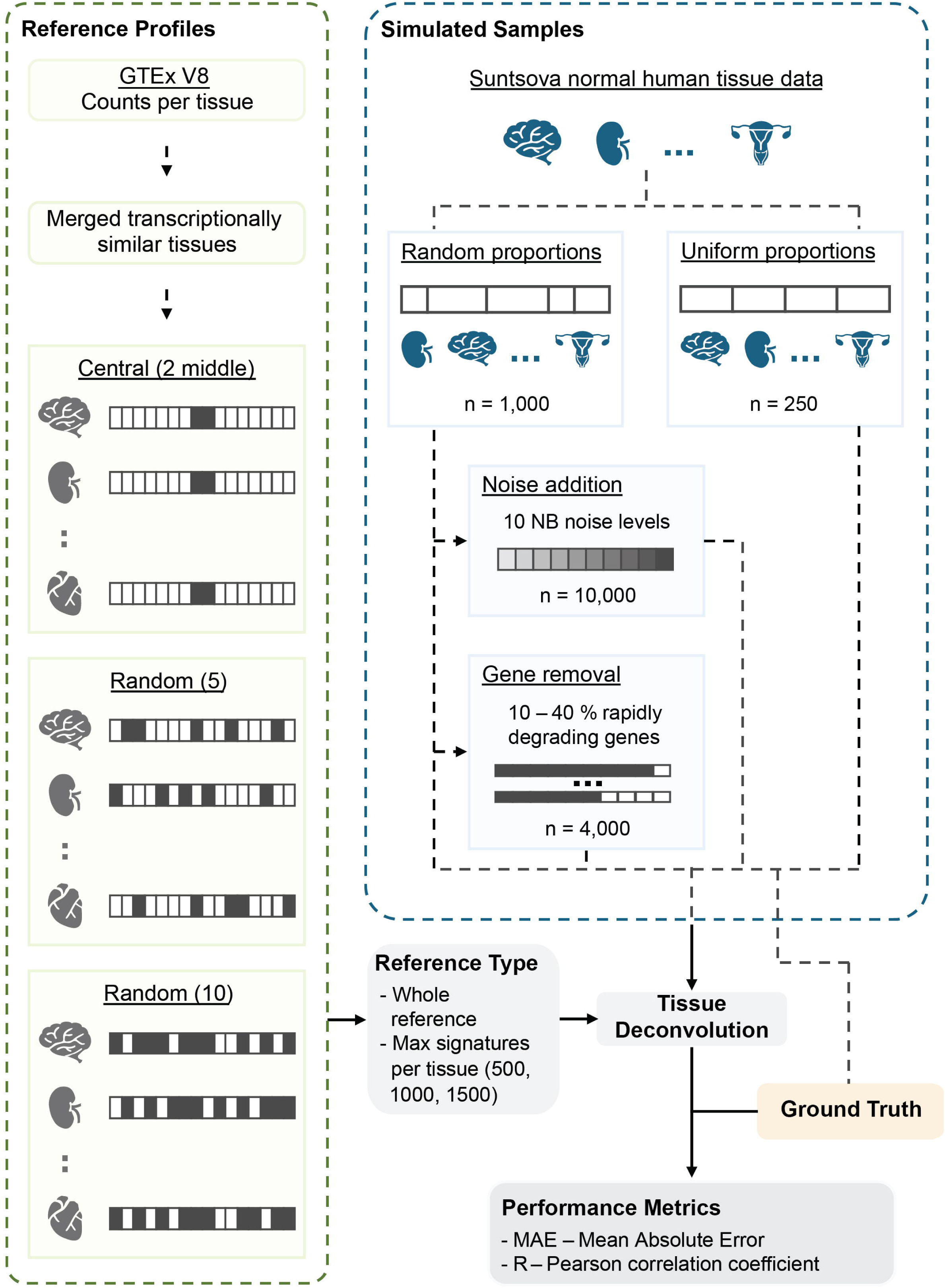
Outline of the tissue-of-origin (TOO) and cell type-of-origin (COO) benchmarking workflow. Overview of the TOO deconvolution framework. GTEx v8 gene read counts per tissue were merged into biological and marker-based tissue groups and used to construct reference expression profiles with either Central (two values closest to the median per tissue), Random-5 (five randomly selected samples per tissue), or Random-10 (ten randomly selected samples per tissue) compositions. These reference profiles were used to generate the required reference input for each deconvolution method. Simulated tissue mixtures were generated by combining tissue data from the same patient using random (n = 1,000) or uniform (n = 250) proportions. Randomly proportioned mixtures were further perturbed by the addition of ten levels of negative binomial noise (n = 10,000) and by removal of deciles of rapidly degrading genes (n = 4,000). Simulated samples and reference profiles were applied to all TOO deconvolution approaches, and performance was evaluated against known ground-truth proportions using mean absolute error (MAE) and Pearson correlation (r).

**Supplementary Figure S2:**
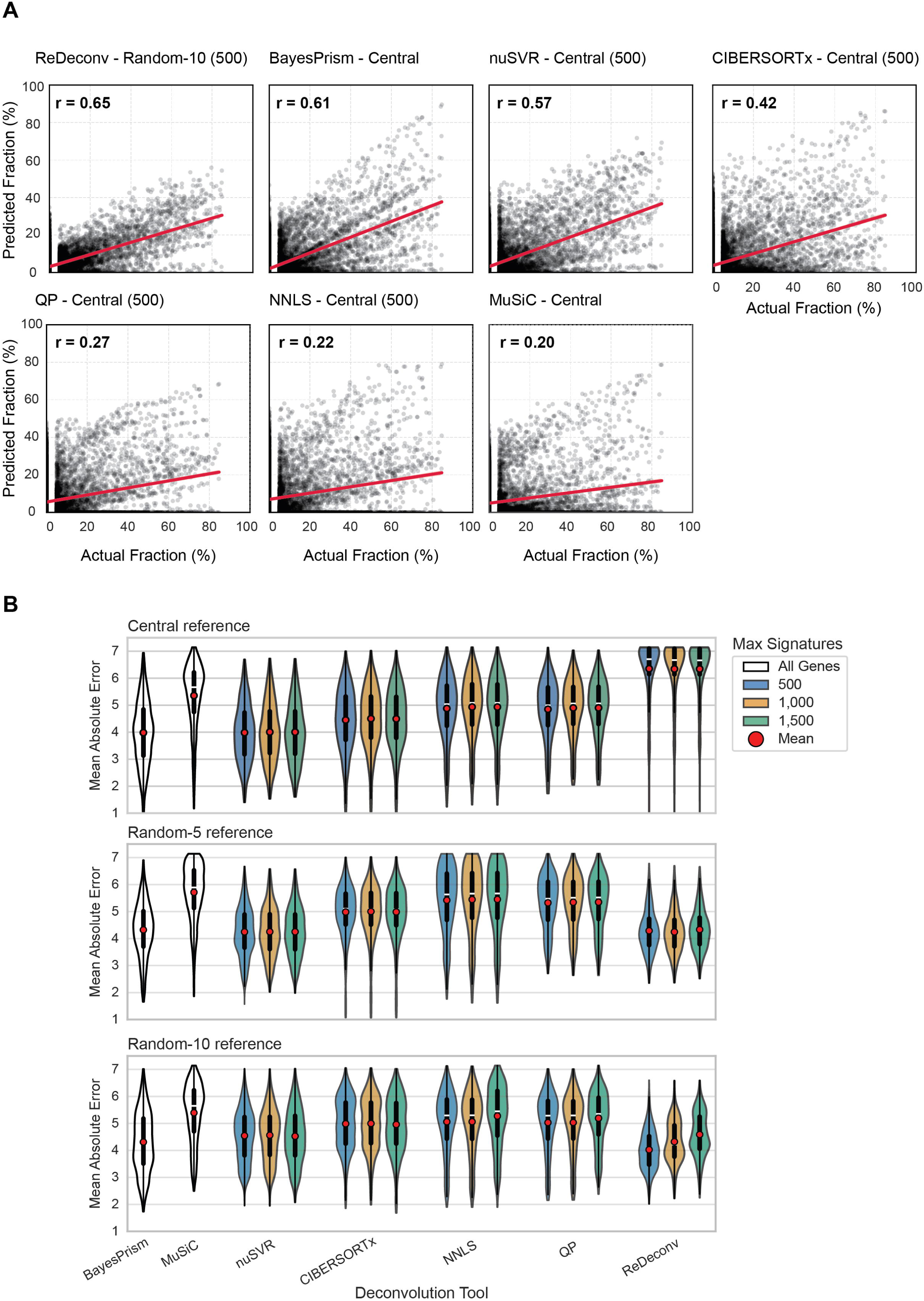
Performance of TOO deconvolution using 1,000 simulated cfRNA mixtures with random proportions. **(A)** Scatterplots showing predicted versus true tissue fractions across 1,000 simulated tissue cfRNA mixtures with random proportions for the best-performing reference configuration and max number of signatures (where applicable) per deconvolution method, defined as the configuration with the highest Pearson correlation (r). **(B)** Distribution of mean absolute error (MAE) across all deconvolution methods and reference construction strategies. Violin plots show MAE distributions for reference matrices constructed using Central, Random-5, or Random-10 tissue sampling approaches, with either all genes or restricted sets of tissue-specific signature genes (500, 1,000, or 1,500). Red points and white horizontal lines inside the violin indicate mean and median MAE values for each method and reference configuration, respectively.

**Supplementary Figure S3:**
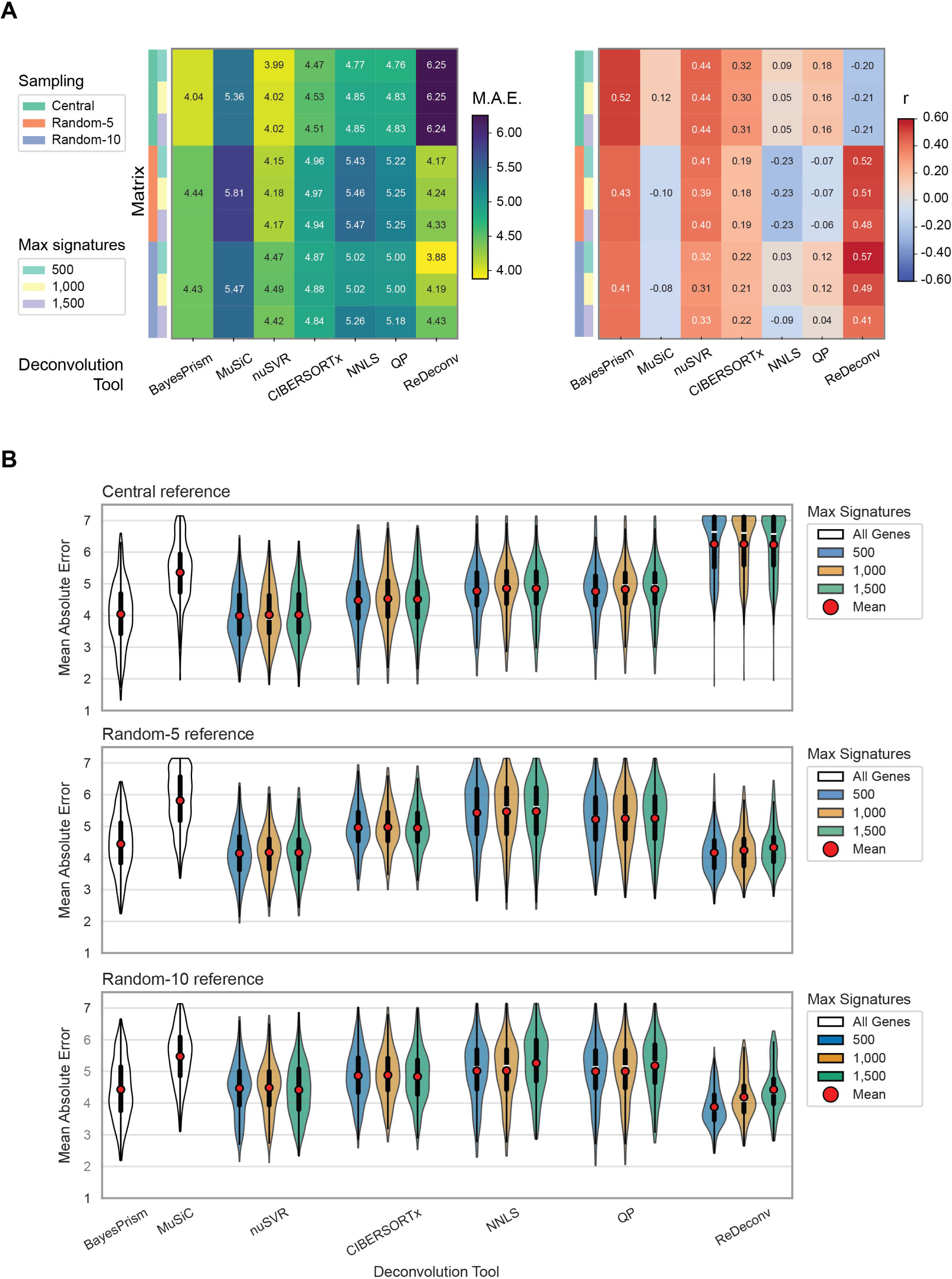
Performance of TOO deconvolution using 250 simulated cfRNA mixtures with uniform proportions. **(A)** Performance of TOO deconvolution for seven methods and nine reference configurations, summarised as heatmaps of (left) mean absolute error (MAE) and (right) Pearson correlation (r) between inferred and true tissue proportions. **(B)** Distribution of MAE across all deconvolution methods and reference construction strategies. Violin plots show MAE distributions for reference matrices constructed using Central, Random-5, or Random-10 tissue sampling approaches, with either all genes or restricted sets of tissue-specific signature genes (500, 1,000, or 1,500). Red points and white horizontal lines inside the violin indicate mean and median MAE values for each method and reference configuration, respectively.

**Supplementary Figure S4:**
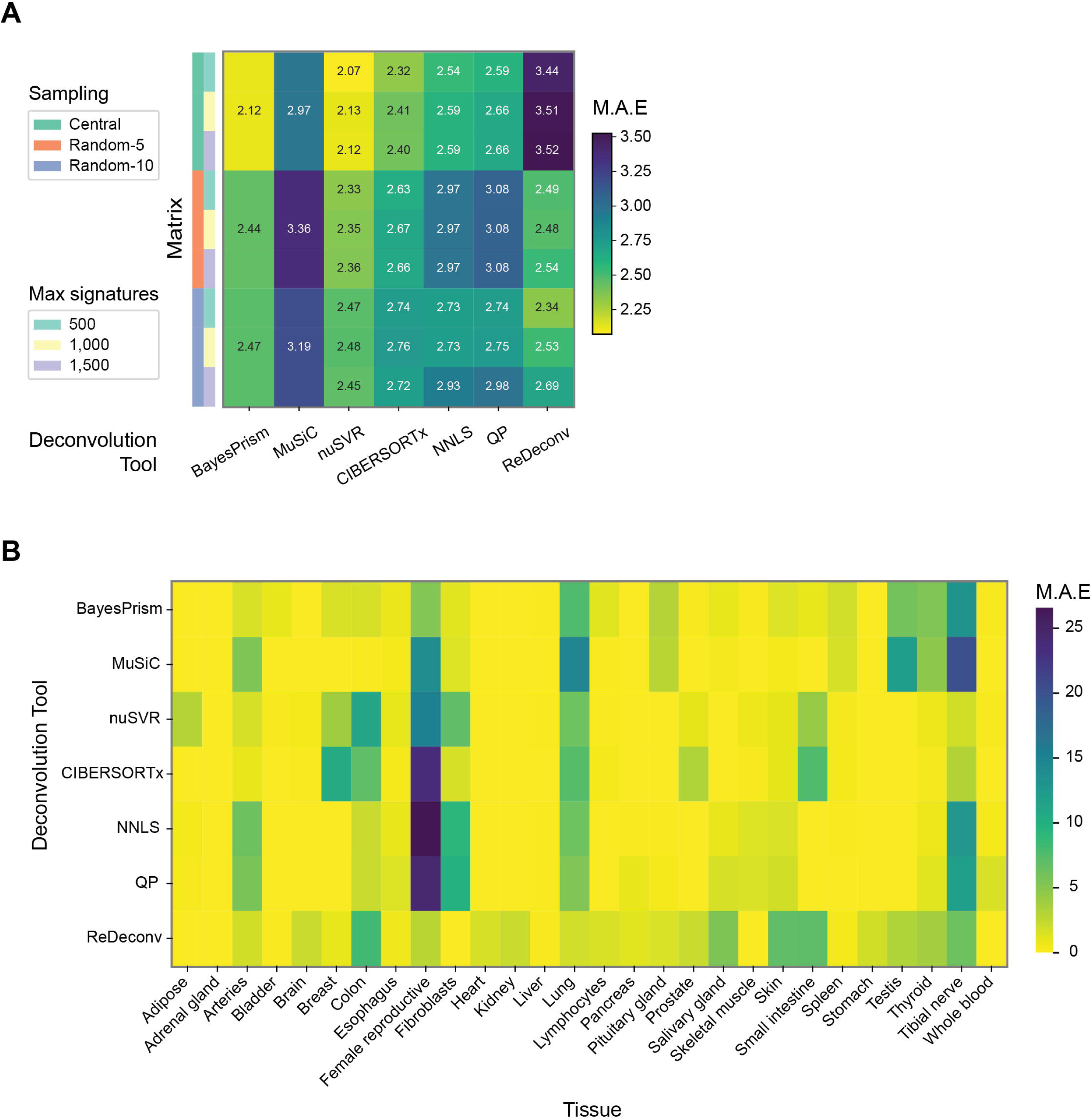
Tissue spillover of TOO deconvolution when tissues are absent from simulated mixtures. **(A)** Mean absolute error (MAE) of tissues absent from 1,000 simulated cfRNA mixtures with random proportions, shown across seven deconvolution methods and nine reference configurations. Spillover error reflects the inferred contribution assigned to tissues not represented in the ground-truth mixture. **(B)** Heatmap showing per-tissue spillover MAE for tissues absent from the simulated mixtures across deconvolution methods, highlighting tissue-specific patterns of misassignment.

**Supplementary Figure S5:**
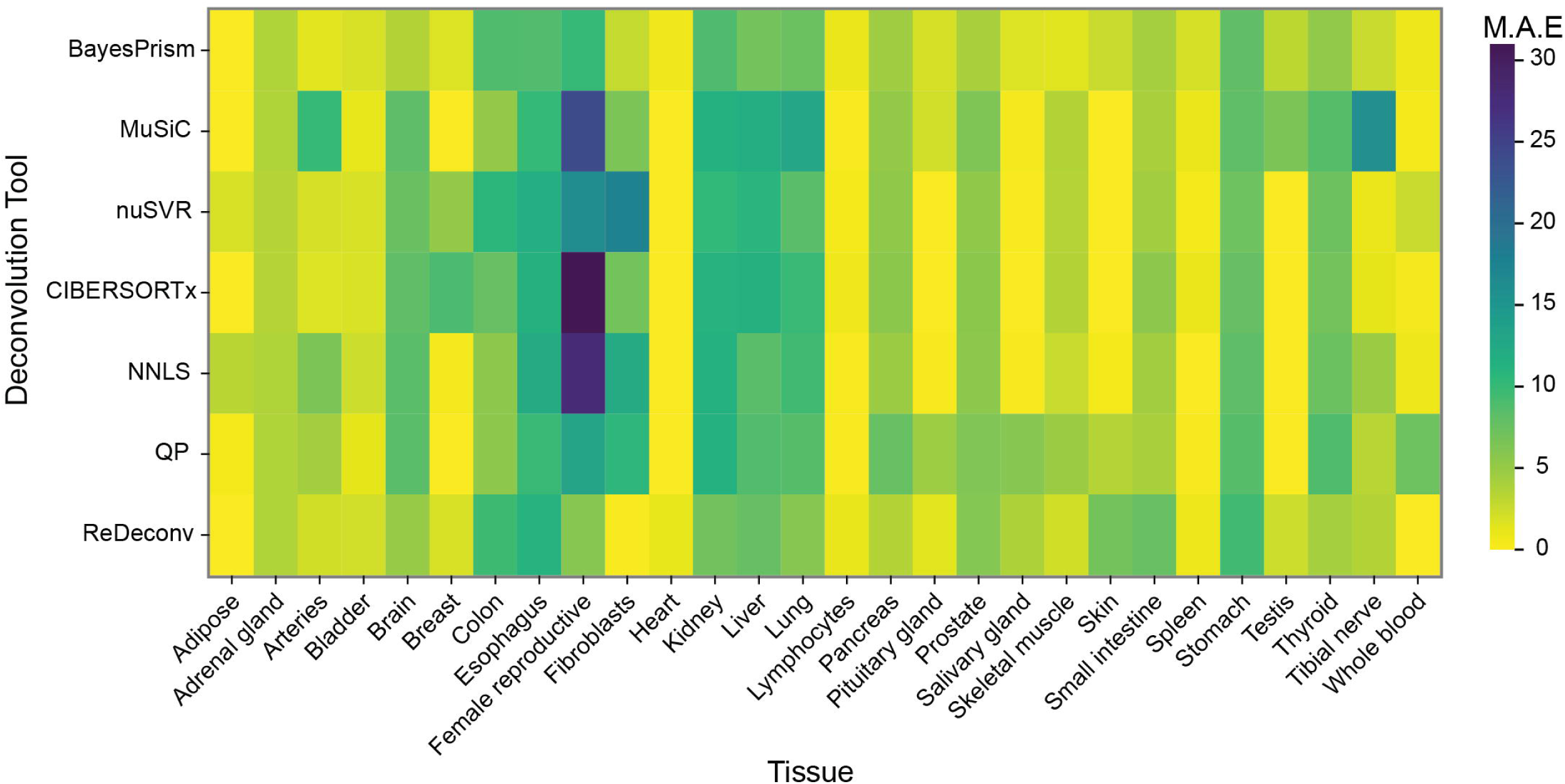
Per-tissue performance of tissue-of-origin (TOO) deconvolution under noise perturbation. Heatmap showing per-tissue mean absolute error (MAE) for 1,000 simulated cfRNA mixtures with random proportions at the highest level of negative binomial noise, across seven deconvolution methods. Results are shown using the best-performing reference configuration for each method, defined by the highest Pearson correlation (r) and lowest mean absolute error (MAE) under baseline (noise-free) conditions.

**Supplementary Figure S6:**
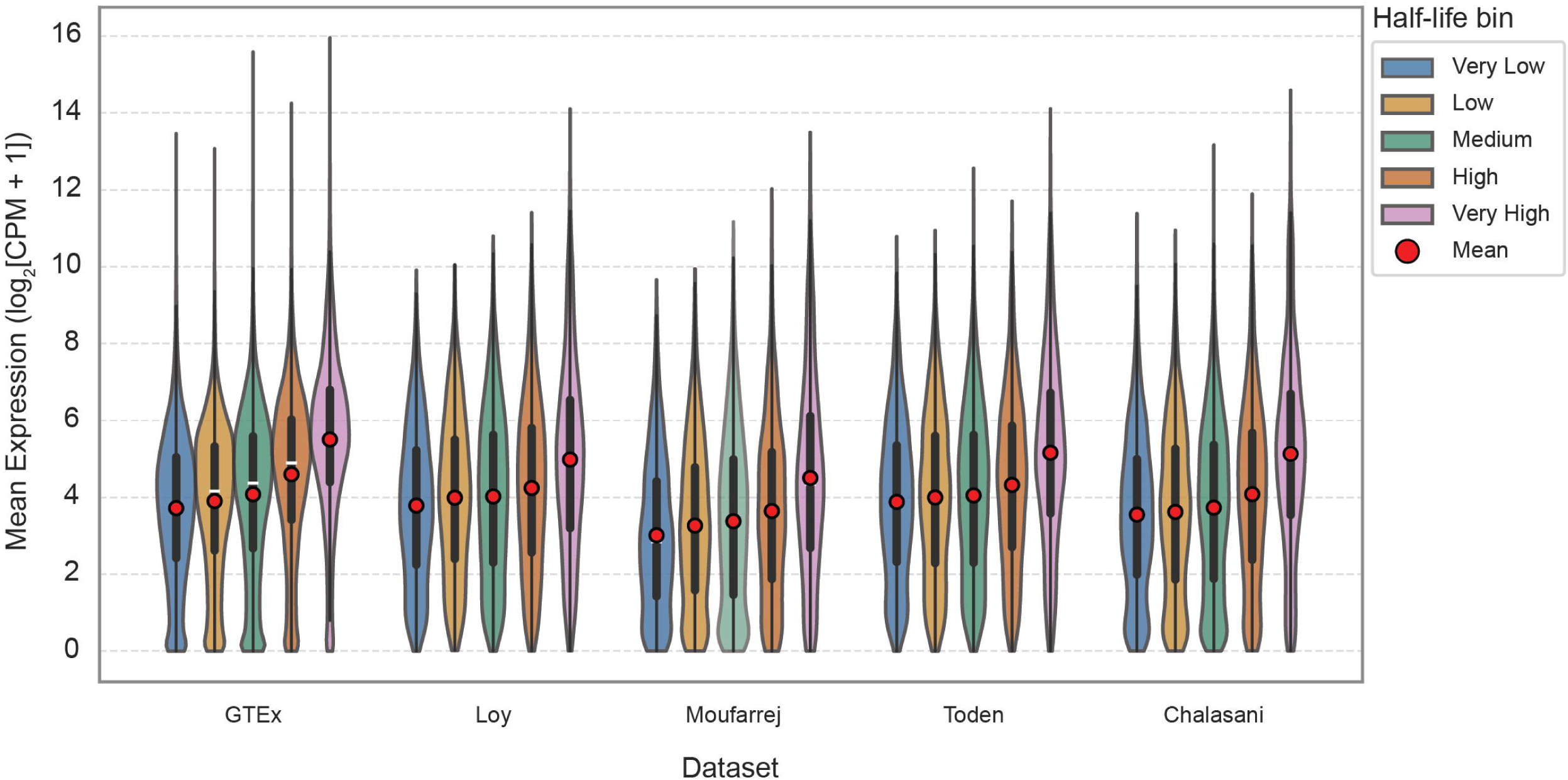
Gene expression stratified by transcript half-life across cfRNA and bulk RNA-seq datasets. Violin plots showing the distribution of gene expression (log₂ mean expression) stratified by transcript half-life bins (very low, low, medium, high, very high) across four cfRNA datasets and GTEx v8 bulk tissue gene read counts. Red points indicate the mean expression within each half-life bin for each dataset, and white horizontal lines within the violins denote the corresponding medians. A monotonic increase in expression across ordered half-life bins was observed in all datasets (Jonckheere–Terpstra trend test, p ≪ 0.001).

**Supplementary Figure S7:**
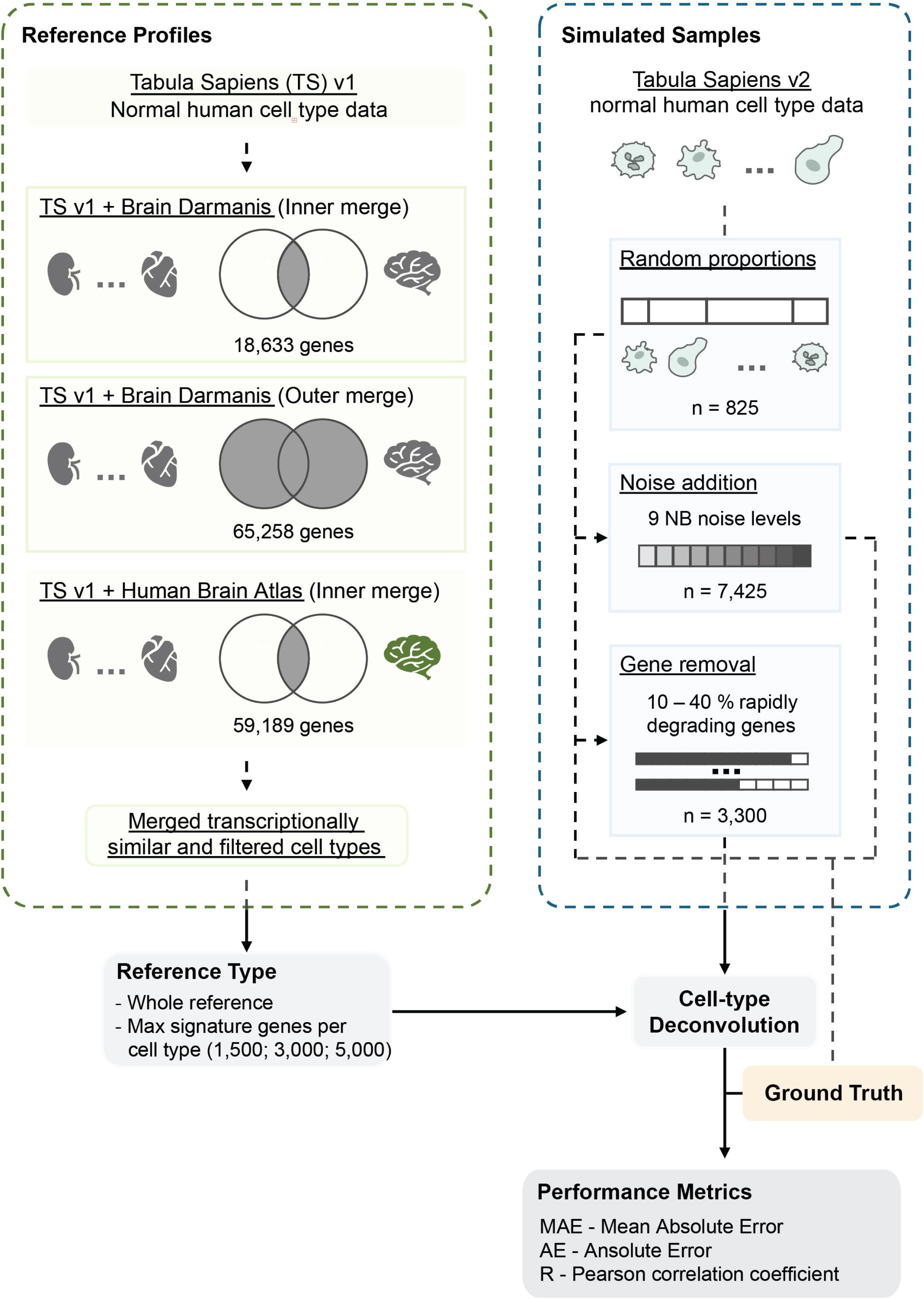
Outline of the cell type-of-origin (COO) benchmarking workflow. Overview of the COO deconvolution framework. Droplet-based single-cell RNA-seq data from Tabula Sapiens (TS) v1 human cell-type data were augmented with augmented with brain-specific cell-type information from the Darmanis Brain dataset (inner and outer merges) and the Human Brain Atlas (inner merge only). These datasets were filtered and transcriptionally similar cell types were merged to generate the reference inputs required for each deconvolution method. Simulated samples (pseudo-bulks) were generated using Tabula Sapiens v2 by combining cell types from the same patients with random proportions (n = 825). These samples were further perturbed by the addition of nine levels of negative binomial noise (n = 7,425) and by removal of deciles of rapidly degrading genes (n = 3,300). Simulated samples and reference profiles were applied to all COO deconvolution approaches, and performance was evaluated against known ground-truth proportions using mean absolute error (MAE), absolute error (AE), and Pearson correlation (r).

**Supplementary Figure S8:**
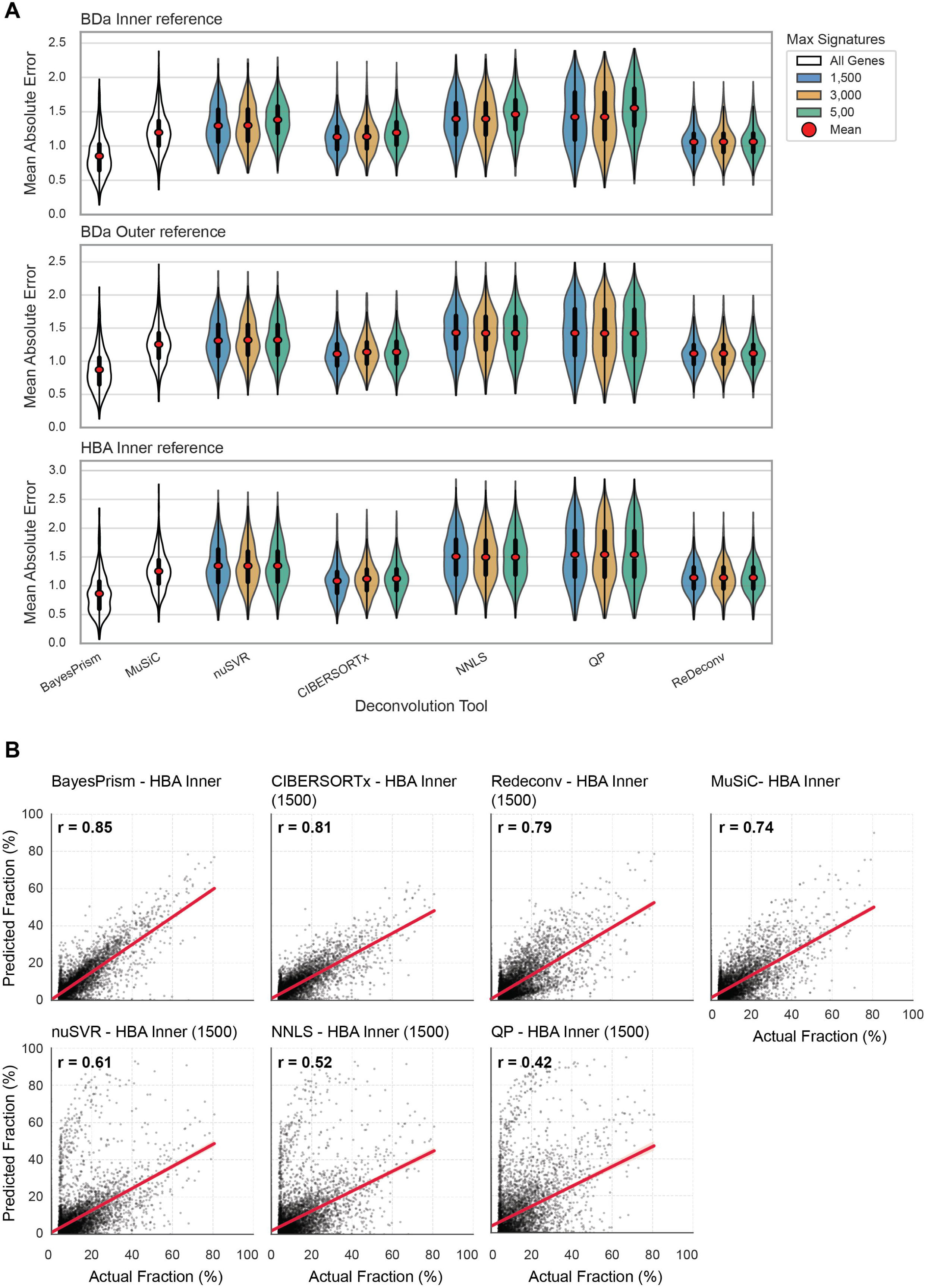
Performance of cell type-of-origin (COO) deconvolution using 825 simulated cfRNA mixtures with random proportions. **(A)** Distribution of mean absolute error (MAE) across all deconvolution methods and reference construction strategies. Violin plots show MAE distributions for reference matrices constructed using Tabula Sapiens v1 augmented with brain single-cell data via BDa Inner, BDa Outer, or HBA Inner gene set merging strategies, using either all genes or restricted sets of cell-type signature genes (1,500, 3,000, or 5,000). Red points and white horizontal lines inside the violin indicate mean and median MAE values for each deconvolution method and reference configuration, respectively. **(B)** Scatterplots showing predicted versus true cell type proportions for the HBA Inner reference configuration and maximum number of signatures (where applicable) per deconvolution method, defined as the configuration yielding the highest Pearson correlation (r).

**Supplementary Figure S9:**
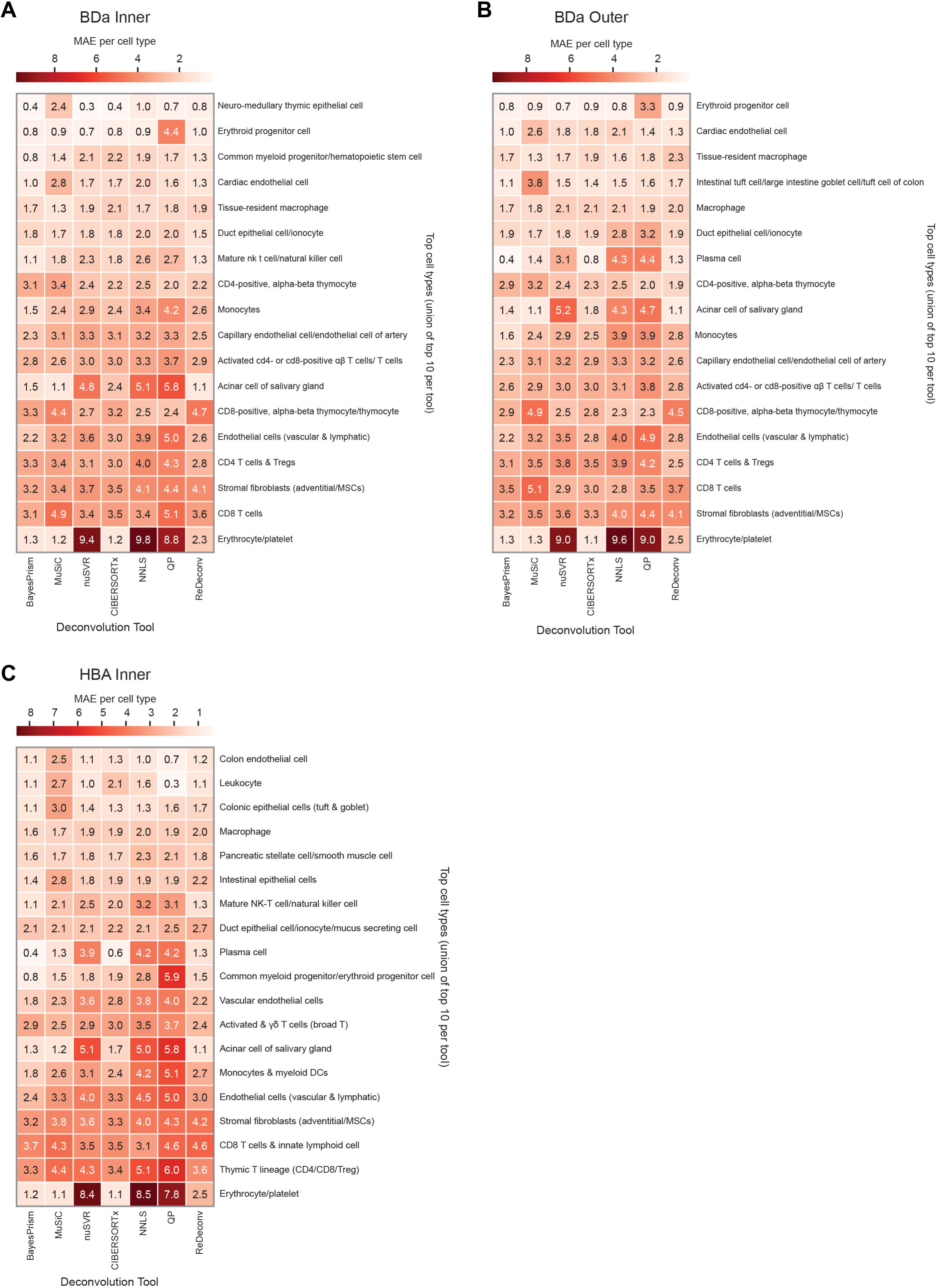
Per–cell-type performance of COO deconvolution in simulated cfRNA mixtures. Mean absolute error (MAE) per cell type for the top 10 cell types with the highest MAE (union across deconvolution tools), evaluated using 825 simulated cfRNA mixtures with random proportions. Heatmaps show results for the best-performing reference configuration of each deconvolution tool using (A) BDa Inner, (B) BDa Outer, or (C) HBA Inner gene-set merging strategies. Values represent mean MAE per cell type, illustrating cell types most prone to spillover across methods. Cell-type labels reflect harmonised groupings derived from merged fine-grained annotations (see **Supplementary Table S2** for full relabelling scheme).

**Supplementary Figure S10:**
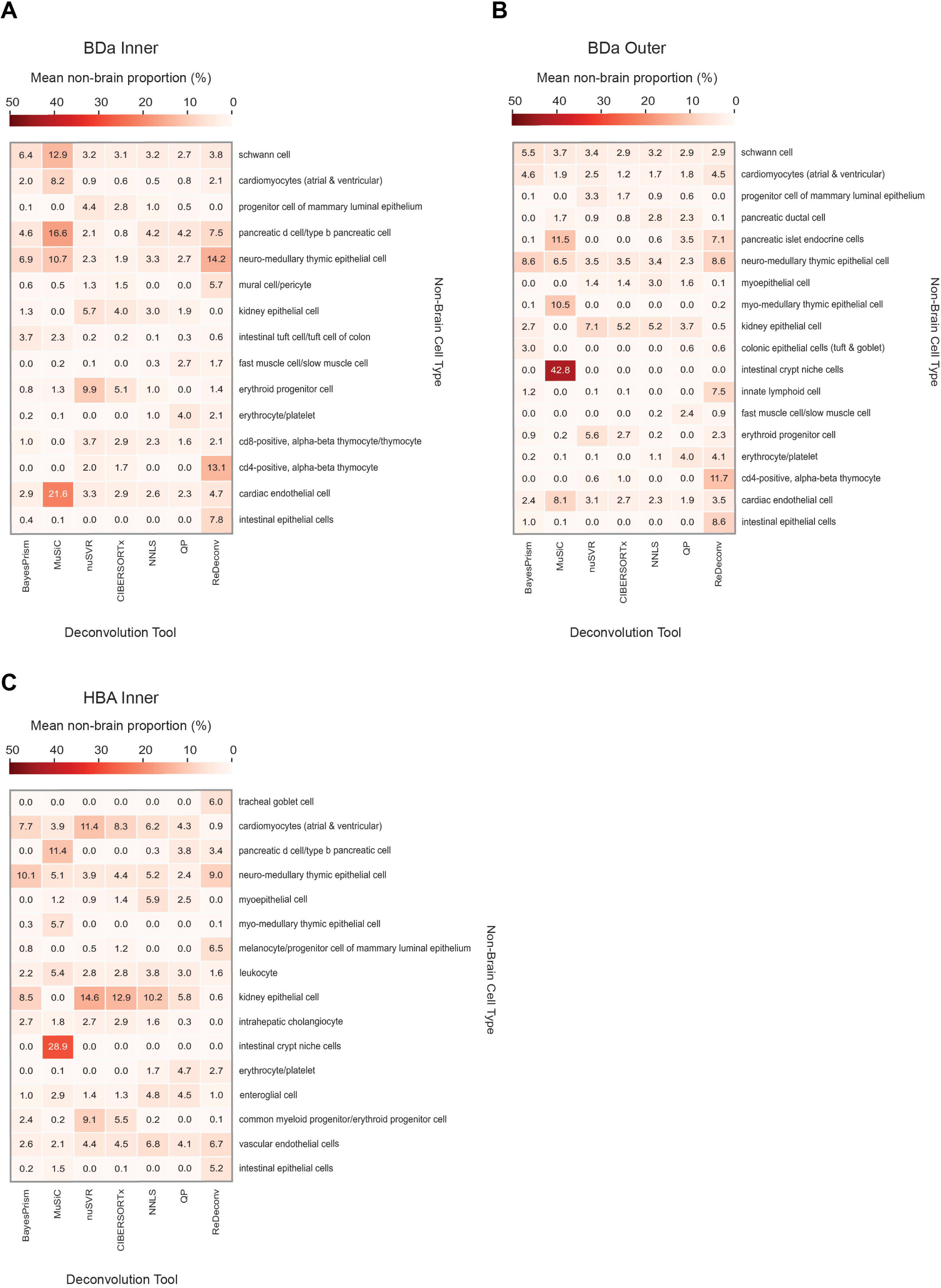
Proportion of non-brain cell types detected in bulk brain samples. Mean proportions (%) of the most abundant non-brain cell types inferred from deconvolution of 200 bulk brain samples from the GTEx v8 dataset. Heatmaps show results for the best-performing reference configuration of each deconvolution tool using **(A)** BDa Inner, **(B)** BDa Outer, or **(C)** HBA Inner gene-set merging strategies. Cell-type labels represent harmonised groupings derived from merged fine-grained annotations across reference datasets (see **Supplementary Table S2** for full relabelling scheme).

**Supplementary Figure S11:**
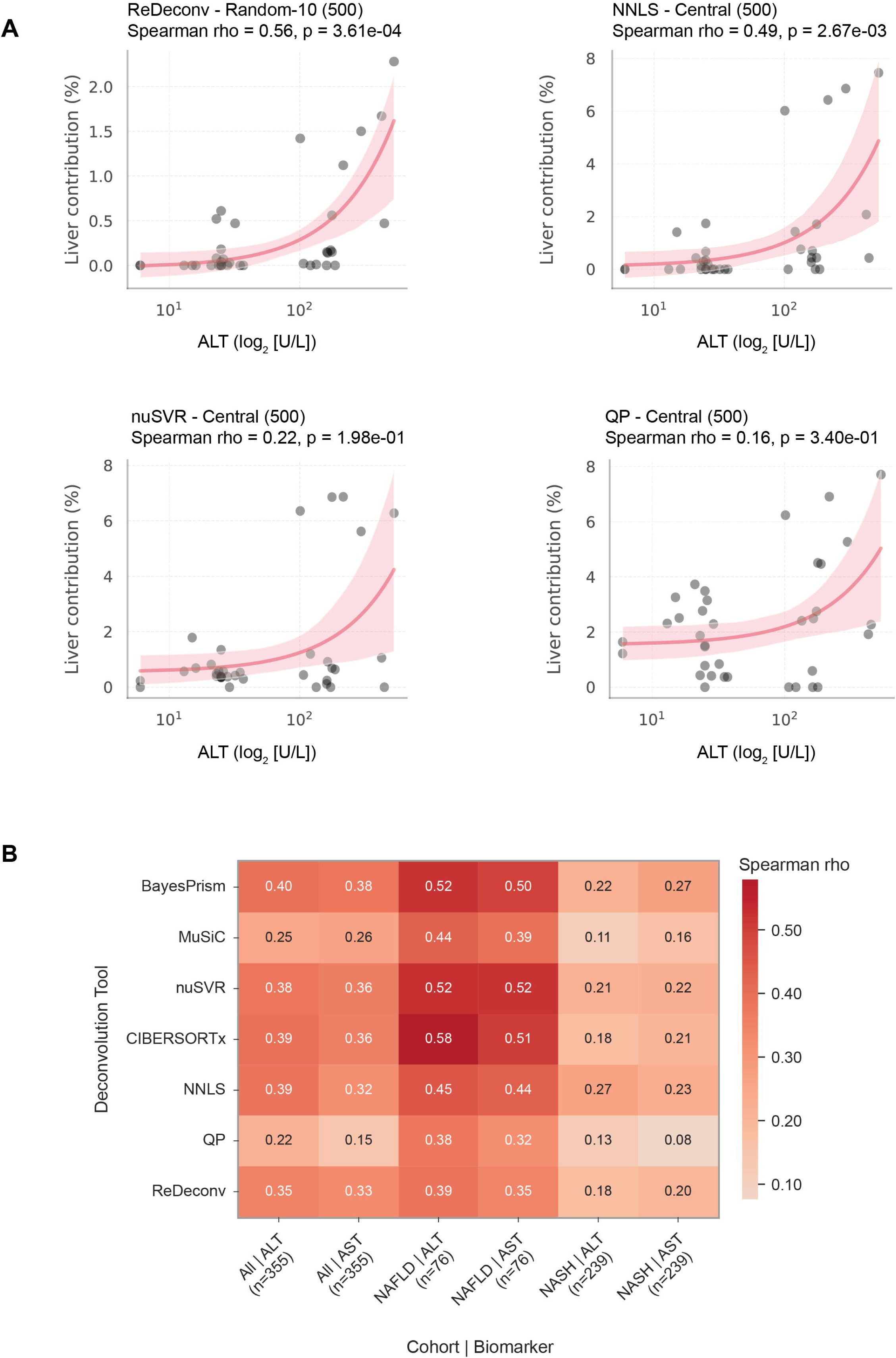
Correlation of TOO deconvolution results with biochemical markers of liver injury. **(A)** Scatterplots of the indicated deconvolution methods illustrating the association between inferred liver contribution and alanine aminotransferase (ALT) levels in a paediatric acute cohort. Spearman rank correlation coefficients are indicated, and fitted trends are shown with shaded confidence intervals. The top-performing methods are shown in the main figure (Fig. 6) The reference matrix and maximum number of signatures used for each deconvolution method are annotated where relevant. **(B)** Heatmap summarising Spearman rank correlations between inferred liver contribution and ALT or aspartate aminotransferase (AST) levels across all deconvolution methods in a chronic liver disease cohort, evaluated in all diseased samples, non-alcoholic fatty liver disease (NAFLD)-only samples, and non-alcoholic steatohepatitis (NASH)-only samples. Deconvolutions were performed using the best-performing reference configuration per method.

**Supplementary Figure S12:**
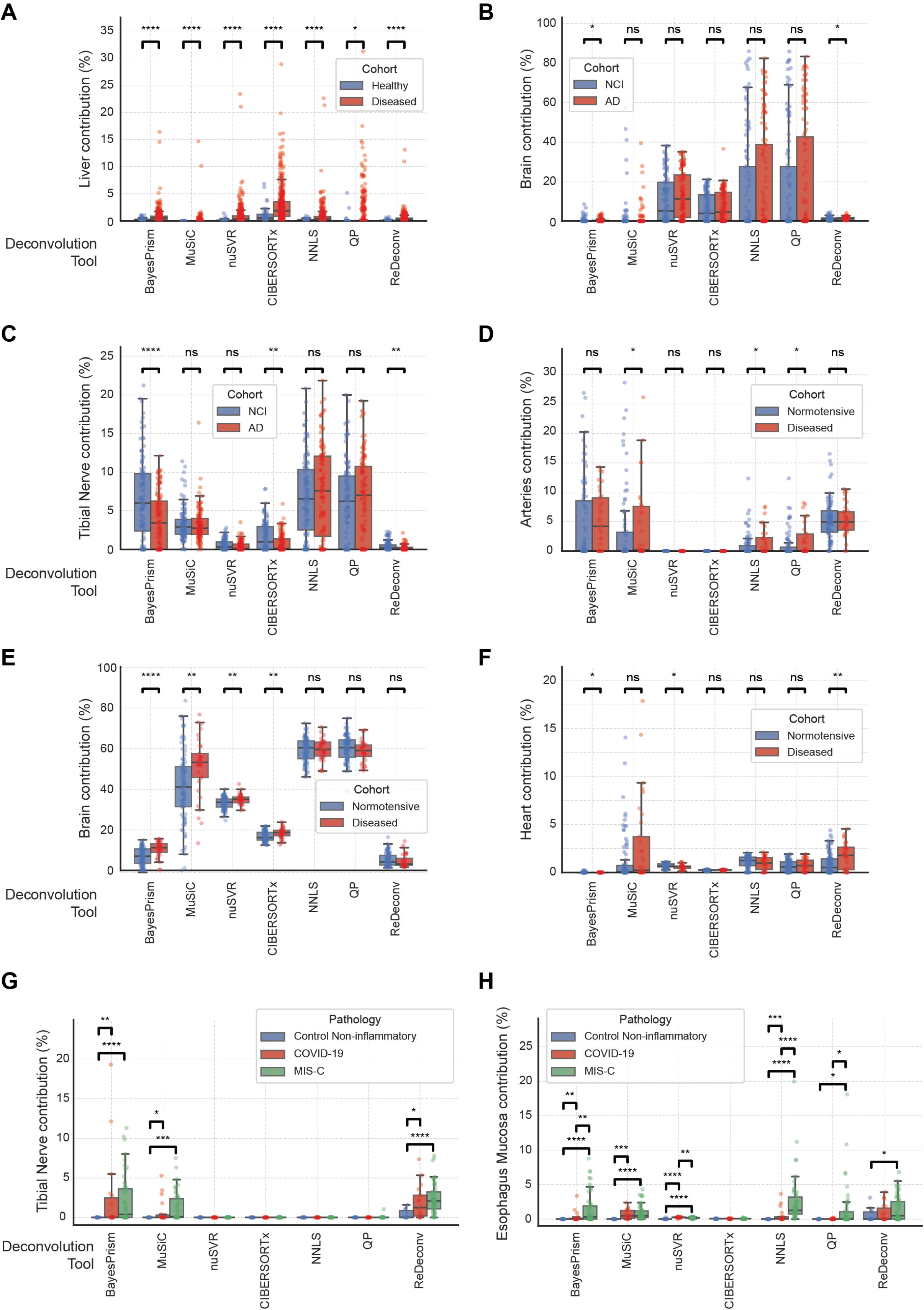
Comparison of inferred tissue contributions across published cfRNA cohorts. **(A)** Comparison of inferred liver contribution between healthy controls and patients with non-alcoholic fatty liver disease or non-alcoholic steatohepatitis (NAFLD/NASH). **(B–C)** Comparison of inferred brain (B) and tibial nerve (C) contributions between Alzheimer’s disease (AD) patients and non–cognitively impaired (NCI) controls. **(D–F)** Comparison of inferred artery (D), brain (E), and heart (F) contributions between normotensive pregnancies and pregnancies complicated by pre-eclampsia or severe pre-eclampsia. **(G–H)** Comparison of inferred tibial nerve (G) and oesophagus mucosa (H) contributions between non-inflammatory controls, patients with COVID-19, and patients with multisystem inflammatory syndrome in children (MIS-C). Distributions of inferred contributions are shown across deconvolution methods with individual samples overlaid. Statistical significance was assessed using two-sided Mann–Whitney U tests; p < 0.05 (*), p < 0.01 (**), p < 0.001 (***), and p < 0.0001 (****).

**Supplementary Figure S13:**
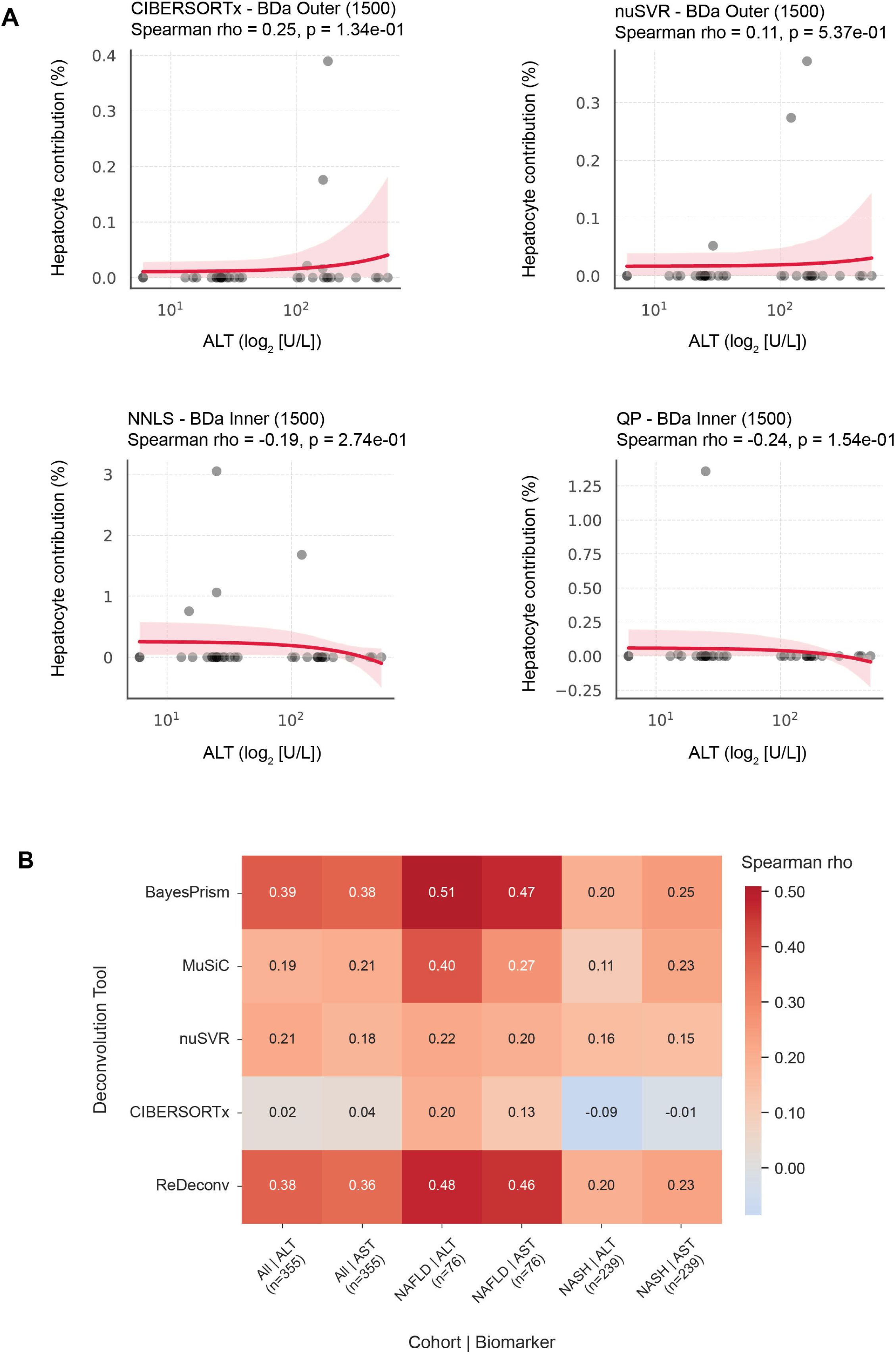
Correlation of COO deconvolution results with biochemical markers of liver injury. **(A)** Scatterplots of the remaining deconvolution methods illustrating the association between inferred hepatocyte contribution and alanine aminotransferase (ALT) levels in a paediatric acute cohort. Spearman rank correlation coefficients are indicated, and fitted trends are shown with shaded confidence intervals. The top-performing methods are shown in the main figure (Fig. 7). Method-specific reference matrices and signature limits are indicated where applicable. **(B)** Heatmap summarising Spearman rank correlations between inferred hepatocyte contribution and ALT or aspartate aminotransferase (AST) levels across all deconvolution methods in a chronic liver disease cohort, evaluated in all samples, non-alcoholic fatty liver disease (NAFLD)-only samples, and non-alcoholic steatohepatitis (NASH)-only samples. Only methods with non-zero inferred hepatocyte signal are shown. All deconvolutions were performed using the best-performing reference configuration per method.

**Supplementary Figure S14:**
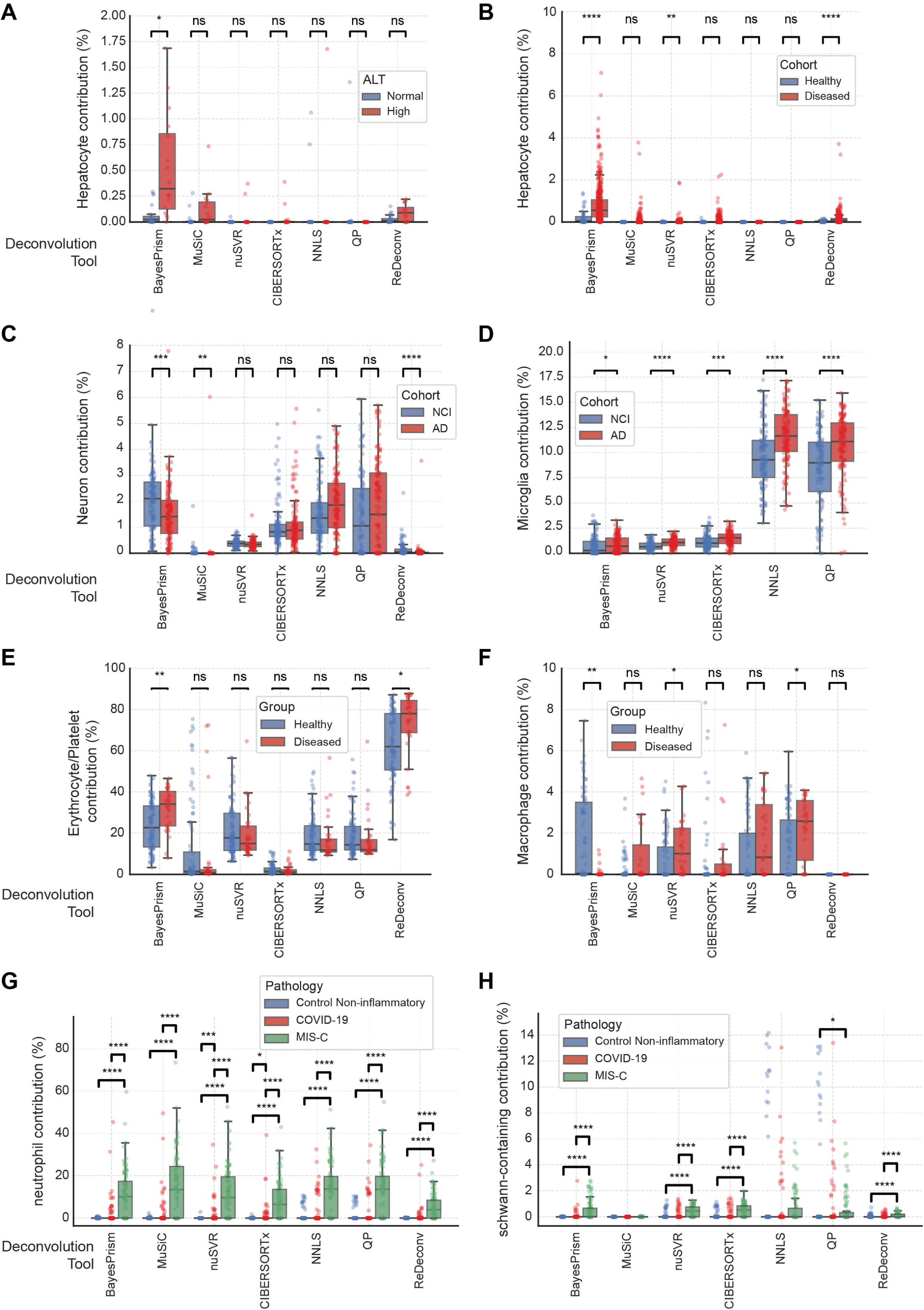
Comparison of inferred cell type contributions across published cfRNA cohorts. **(A)** Comparison of inferred hepatocyte contribution between patients with high and normal ALT levels in a paediatric acute cohort. **(B)** Comparison of inferred hepatocyte contribution between healthy controls and patients with non-alcoholic fatty liver disease or non-alcoholic steatohepatitis (NAFLD/NASH). **(C–D)** Comparison of inferred neuronal (C) and microglia-containing (D) group contribution between Alzheimer’s disease (AD) patients and non–cognitively impaired (NCI) controls. **(E–F)** Comparison of inferred erythrocyte/platelet (E) and macrophage (F) contribution between normotensive pregnancies and pregnancies complicated by pre-eclampsia or severe pre-eclampsia. **(G–H)** Comparison of inferred neutrophil (G) and Schwann cell-containing group (H) contributions between non-inflammatory controls, patients with COVID-19, and patients with multisystem inflammatory syndrome in children (MIS-C). Statistical significance was assessed using two-sided Mann–Whitney U tests with Benjamini–Hochberg false discovery rate correction applied within each deconvolution method; p < 0.05 (*), p < 0.01 (**), p < 0.001 (***), and p < 0.0001 (****).

**Supplementary Figure S15:**
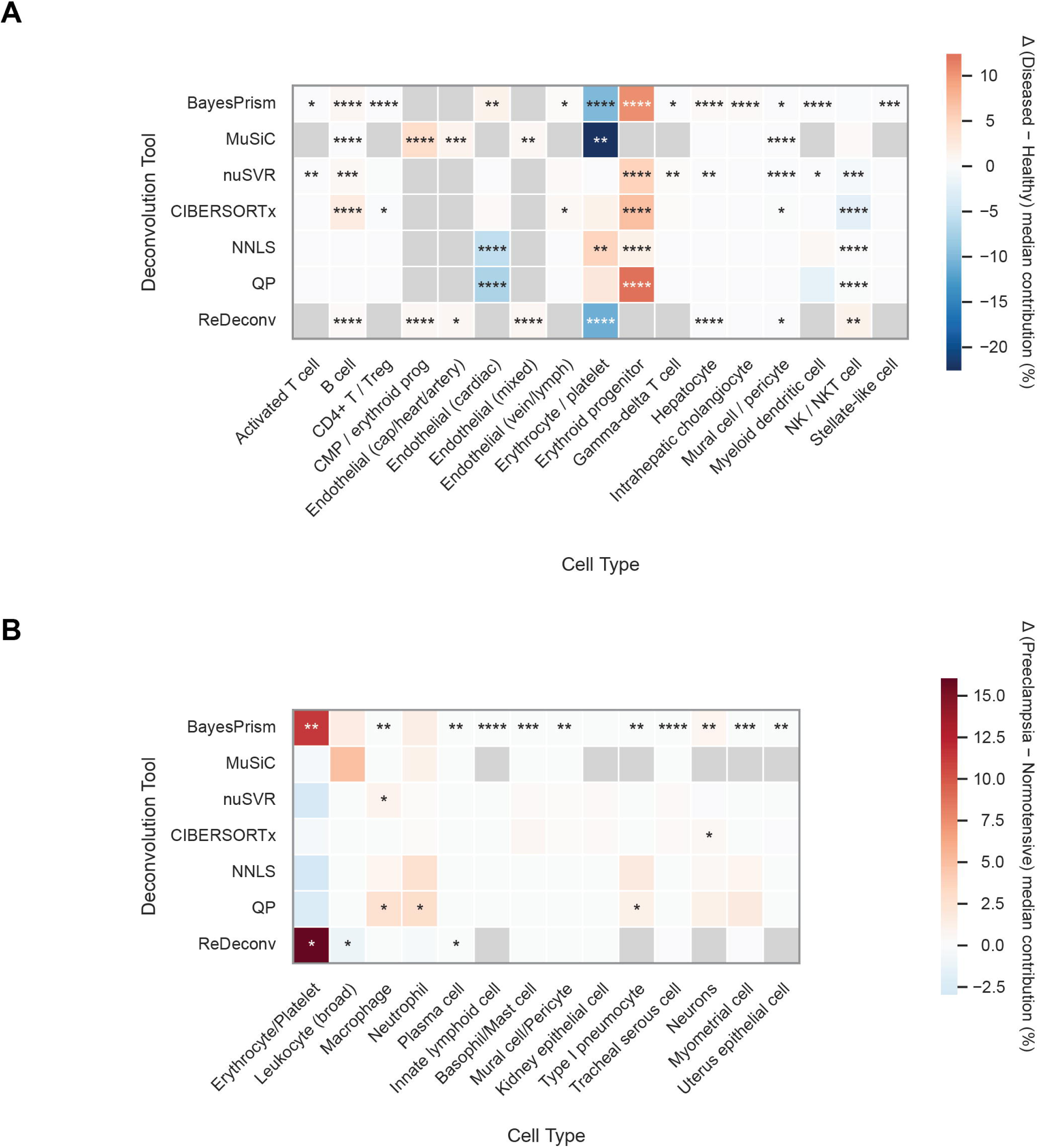
Disease-associated differences in inferred cell-type contributions from COO deconvolution. **(A)** Difference in median inferred cell-type contributions between patients with non-alcoholic fatty liver disease or non-alcoholic steatohepatitis (NAFLD/NASH) and healthy controls. Values represent the difference (Δ) in cell-type contribution between disease and control (median contribution in the disease group minus the median contribution in healthy controls) for each deconvolution method and cell type. **(B)** Difference (Δ) in median inferred cell-type contributions between pregnancies with pre-eclampsia or severe pre-eclampsia and normotensive pregnancies. Values represent the median contribution in diseased pregnancies minus that in normotensive controls for each deconvolution method and cell type. Grey boxes indicate cell-type categories absent from the reference matrix used by the corresponding deconvolution method. Statistical significance was assessed using two-sided Mann–Whitney U tests with Benjamini–Hochberg false discovery rate correction applied within each deconvolution method; p < 0.05 (*), p < 0.01 (**), p < 0.001 (***), and p < 0.0001 (****).

